# The Human RNA-Binding Proteome and Its Dynamics During Arsenite-Induced Translational Arrest

**DOI:** 10.1101/329995

**Authors:** Jakob Trendel, Thomas Schwarzl, Ananth Prakash, Alex Bateman, Matthias W. Hentze, Jeroen Krijgsveld

## Abstract

Proteins and RNA functionally and physically intersect in multiple biological processes, however, currently no universal method is available to purify protein-RNA complexes. Here we introduce XRNAX, a method for the generic purification of protein-crosslinked RNA, and demonstrate its versatility to study the composition and dynamics of protein-RNA interactions by various transcriptomic and proteomic approaches. We show that XRNAX captures all RNA biotypes, and use this to characterize the sub-proteomes that interact with coding and non-coding RNAs (ncRNAs), and to identify hundreds of protein-RNA interfaces. Exploiting the quantitative nature of XRNAX, we observe drastic remodeling of the RNA-bound proteome during arsenite-induced stress, distinct from autophagy-induced changes in the total proteome. In addition, we combine XRNAX with CLIP-seq to validate the interaction of ncRNA with Lamin B and EXOSC2. Thus, XRNAX is a resourceful approach to study structural and compositional aspects of protein-RNA interactions to address fundamental questions in RNA-biology.

## Introduction

Cellular proteins and RNA intimately interact in intricate ways to regulate a wide range of processes that are essential for cells to survive, replicate, and adapt to environmental changes. For instance, proteins accompany messenger RNA (mRNA) throughout its lifetime from transcription to decay, deploying specialized complexes for RNA splicing, capping, translation and localization (for review see (Müller-McNicoll and Neugebauer, 2013)). Indeed, in molecular machines like the ribosome proteins and various types of RNAs (i.e. ribosomal RNA (rRNA), transfer RNA (tRNA) and mRNA) converge for the genesis of new proteins. Given the central role of such processes in the flow of genetic information, multiple techniques have been developed to characterize protein-RNA interfaces, primarily driven by sequencing technologies. This includes crosslinking and immunoprecipitation followed by sequencing (CLIP-seq) (Ule et al., 2003) and its many derivatives (for review see (Lee and Ule, 2018)) to identify RNA that interact with a protein of interest, as well as ribosome profiling(Ingolia et al., 2009) to identify RNA bound to ribosomes in the act of translation. More recently this has been complemented with proteomic technologies to approach protein-RNA interactions from the opposite end and identify proteins that interact with RNA. In particular, we (Beckmann et al., 2015; Castello et al., 2012, 2013; Kwon et al., 2013; Liepelt et al., 2016; Sysoev et al., 2016) and others (Baltz et al., 2012; Matia-González et al., 2015; Mitchell et al., 2012; Wessels et al., 2016) have successfully employed poly(dT) capture for the enrichment of proteins interacting with polyadenylated RNA (i.e. primarily mRNA) in various cell types and organisms. During this procedure, termed RNA interactome capture, proteins interacting with RNA are crosslinked *in vivo* by UV-irradiation, polyadenylated RNA-protein complexes are captured via oligo(dT)-coated beads, and proteins are identified by mass spectrometry (MS). This approach has proven very powerful for identifying novel RNA-binding proteins (Castello et al., 2012), for quantifying changes in protein-mRNA interactions (Sysoev et al., 2016), and for the characterization of RNA-binding sequence features and domains (Castello et al., 2016) (for review see (Hentze et al., 2018)).

Despite this progress, these proteomic methods are limited to only a subset of the transcriptome: since RNA must be polyadenylated for it to be captured, interactome capture ignores >95% of mammalian cellular RNA (per weight) that is non-polyadenylated (Hastie and Bishop, 1976).This includes highly abundant ‘housekeeping’ RNAs such as tRNA, rRNA, small nuclear RNAs (snRNAs), small nucleolar RNAs (snoRNAs), but also a diverse group of other non-coding RNAs (ncRNAs) whose functionalities are only beginning to emerge (for review see (Long et al., 2017; Wang and Chang, 2011)). Like for mRNA, the biogenesis and functionality of these RNA species is determined primarily by their interaction with protein, however currently no universal method is available for the comprehensive purification of protein-RNA complexes from living cells.

Inspired by the perspective that an unbiased, comprehensive characterization of protein-RNA interactions will greatly aid in understanding the functionality of both the RNA and protein involved, we developed a method for the extraction and purification of protein-crosslinked RNA irrespective of RNA biotype. We named this method XRNAX, for ‘protein-crosslinked RNA eXtraction’. Crucially, XRNAX purifies protein-crosslinked RNA as a physical entity that can serve as a universal starting point for multiple downstream applications, both using sequencing and MS. Here, we combined XRNAX with proteomics to derive an integrated draft of the human RNA-binding proteome from three cell lines and report distinctive features that distinguish proteins binding to mRNA or ncRNA. In addition, we located protein-RNA interfaces through ribonucleotide-crosslinked peptides and identified evidence for a novel RNA-binding domain. Subsequently, we quantified the dynamics in protein-RNA interactions following arsenite treatment, and report massive remodeling of the translation-associated and RNA-interacting proteome in human cells. We show that this is mediated by autophagy in a process that is exceptionally fast, eliminating 50% of ribosomal proteins and many other RNA-binding proteins within 30 minutes. Finally, we used XRNAX prior to CLIP-seq, validating LMNB1 as a novel RNA-binder interacting with snoRNAs, and showing that the nuclear translocation of the RNA exosome component EXOSC2 is concomitant with a change in its decoration with RNA. Collectively, we demonstrate that XRNAX is a powerful tool for the discovery of novel RNA-interacting proteins by MS and their validation through CLIP-seq even from the identical sample. We envision that this will be of broad utility to understand fundamental regulatory processes in many cell types and organisms.

## Results

### Organic Phase Separation for the Extraction of Protein-Crosslinked RNA

Since interactome capture is restricted to the identification of proteins that interact with polyadenylated RNA, we sought to design a method to investigate protein-RNA interactions irrespective of the RNA biotype, and that would enable the characterization of the RNA-bound proteome using MS, but also of the protein-bound transcriptome using RNA sequencing.

Since acid guanidinium thiocyanate–phenol–chloroform extraction (in the following called TRIZOL (Chomczynski and Sacchi, 1987)) is a classical method to separate RNA (in the aqueous phase) from protein (in the organic phase), we hypothesized that protein-crosslinked RNA might end up in the insoluble TRIZOL interphase (Figure 1A). We therefore UV-crosslinked MCF7 cells and collected the TRIZOL interphase, washed it to remove free protein and RNA, and DNase-digested to eliminate DNA, yielding an extract containing highly concentrated RNA (>1000 ng/µl) and protein (>0.7 mg/ml) (see STAR-methods for details). Interestingly, when run on an agarose gel this extract showed a band with a distinct upward shift compared to conventional TRIZOL-extracted RNA (Figure 1B, arrows). This band disappeared upon RNase digestion, showing that it contained RNA without contaminating DNA. Similarly it disappeared after proteinase-K digestion, collectively showing that the extract material consisted of protein-crosslinked RNA. In addition, identical results were achieved when applying UV-crosslinking at 365 nm using cells grown in the presence of the photoactivatable nucleotide 4-thiouridine (4-SU) (Figure 1B), showing that this procedure is independent of the type of UV-crosslinking. In conclusion, this indicated that the procedure, consisting of UV-crosslinking followed by TRIZOL extraction and subsequent processing steps of the interphase, specifically purified protein-crosslinked RNA. We termed this method protein-crosslinked RNA eXtraction, or XRNAX (Figure 1A).

**Figure 1:**
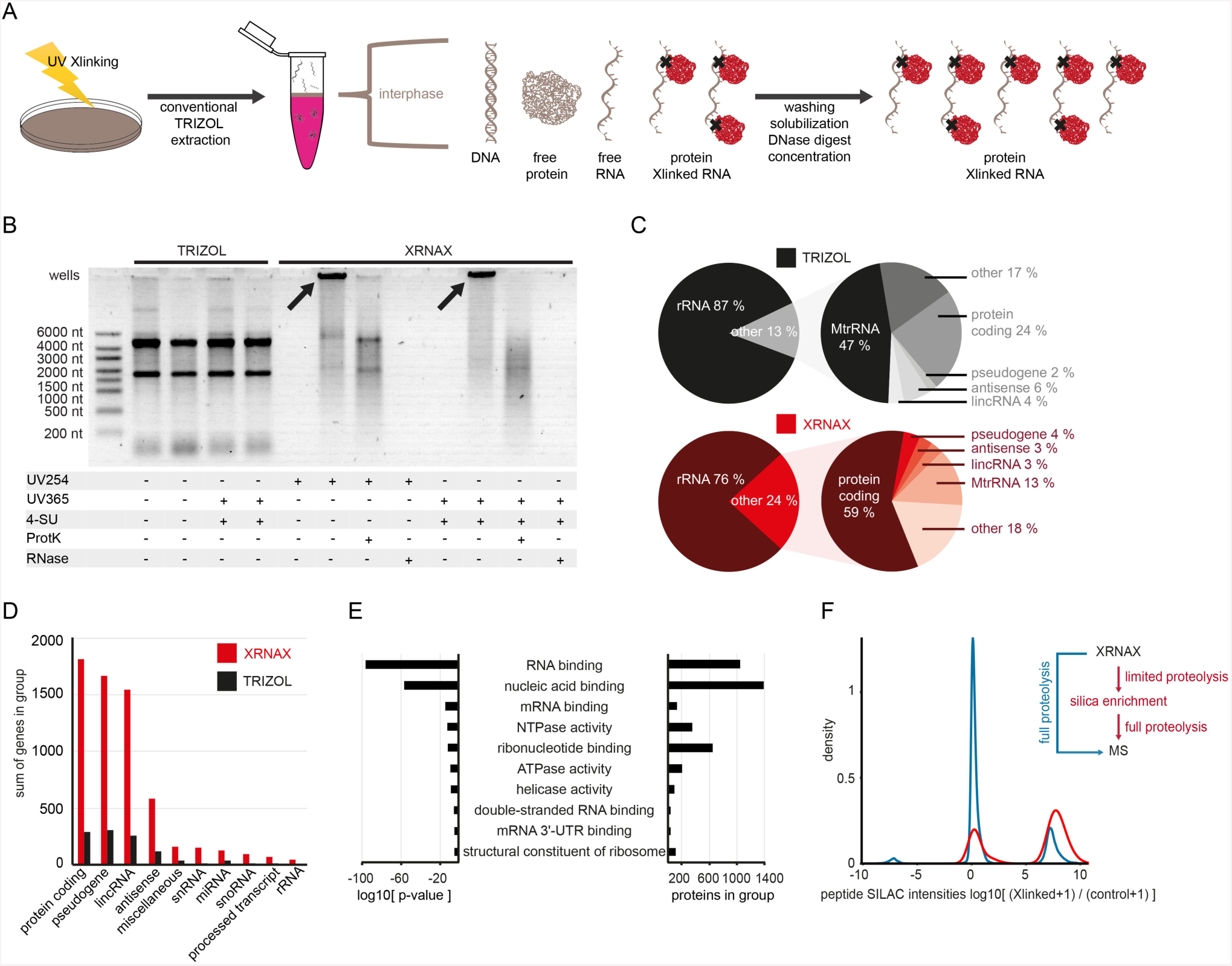
XRNAX extracts protein-crosslinked RNA from UV-crosslinked cells. (A) Experimental scheme of XRNAX. (B) Comparison of classic TRIZOL and XRNAX by agarose gel electrophoresis. MCF7 cells were subjected to crosslinking with UV at 254 nm (UV254) or incubated with 100 µM 4-thiouridine (4-SU) for 16 hours and crosslinked at 365 nm (UV365). RNA was extracted using either conventional TRIZOL or XRNAX and subsequently treated as indicated. ProtK: Digestion with proteinase K. RNase: Digestion with RNase A and I. (C) Pie diagrams comparing RNA composition of TRIZOL extracts (top) in comparison to XRNAX extracts (bottom). RNA from XRNAX extracts or TRIZOL extracts, was sequenced before (left) and after (right) depletion of ribosomal RNA, respectively. Reads for each GENCODE biotype were normalized to the total number of reads in one library. (D) Transcripts exclusively identified after sequencing of either TRIZOL or XRNAX-extracted RNA from MCF7 cells that were grown in the presence of 4-SU. Sum of genes are summarized within the top-10 GENCODE biotypes. (E) GO-enrichment analysis for proteins in XRNAX extracts from MCF7 cells. Displayed are the ten most-enriched terms. (F) Density plot showing SILAC ratios of peptides from MCF7 XRNAX-extracts before (blue) and after silica purification (red). Pseudo-counts were added to display peptides exclusively identified in either SILAC channel.

### XRNAX-Extracts Contain RNA of All Major Biotypes

We next used various transcriptomic and proteomic techniques to characterize the RNA and protein composition of XRNAX extracts. To first obtain a global view on the type and relative distribution of RNA biotypes, we digested the XRNAX extract obtained from MCF7 cells with proteinase K and sequenced its RNA. For comparison we sequenced total RNA extracted from non-crosslinked MCF7 cells using the conventional TRIZOL protocol. In addition, RNA was sequenced before and after depletion of rRNA to display the entire relative composition of each preparation. This showed that XRNAX extracts contained all major RNA biotypes also detected in classical TRIZOL extracted RNA, albeit in different relative amounts (Figure 1C). Specifically, XRNAX extracts contained double the amount of non-rRNA and three times less mitochondrial rRNA, likely reflecting the selective enrichment of protein-bound RNA. Very similar results were obtained when using 4-SU labeling and crosslinking at 365 nm. Interestingly, we found that, at virtually identical sequencing depth, libraries prepared via XRNAX had much better coverage for medium and low abundant transcripts (Figure S1B and C). Effectively, this led to the detection of 6306 transcripts exclusively observed in XRNAX-derived libraries, including ∼1500 protein-coding and ∼1500 lincRNAs (Figure 1D). These results demonstrated that XRNAX-extracts contained all major RNA biotypes.

### XRNAX Extracts Are Enriched In RNA-Binding Proteins UV-Crosslinked to RNA

Next we used MS to profile the protein composition of XRNAX extracts and to validate that these contain protein that is crosslinked to RNA. We therefore subjected MCF7 cells to XRNAX and compared protein composition of the extract to the total proteome obtained from an MCF7 cell lysate. Gene ontology (GO) enrichment analysis revealed various terms related to RNA binding and function as the most significantly enriched and densely populated categories (Figure 1E). Ranking by protein abundance using intensity-based absolute quantification (iBAQ) (Schwanhäusser et al., 2011) showed a shift from histones and cytoskeletal proteins as the most abundant proteins in the total proteome, to HNRNPs, splicing factors and other RNA-binding proteins in XRNAX samples (Figure S1A). Collectively this illustrates that XRNAX strongly enriched for RNA-binding proteins.

To control for unspecific or indirect protein-protein rather than direct protein-RNA interactions we applied stable isotope labeling in cell culture (SILAC) in three validation experiments. First, heavy SILAC-labeled cells were UV-crosslinked and combined with non-crosslinked light SILAC-labeled cells, subjected to XRNAX and the protein content of the extract analyzed by MS. This revealed many peptides that only showed intensity in the heavy SILAC channel, indicating enrichment of RNA-crosslinked peptides (Figure 1F). However, we noted that many non-enriched peptides originated from *bona fide* RNA-binding proteins, suggesting that they were trapped in the TRIZOL interphase even without crosslinking, potentially as a result of the preferred interaction of RNA-binding proteins with other RNA-binding proteins (Brannan et al., 2016).

To solve this, we designed a denaturing silica-based cleanup procedure downstream of XRNAX. Since silica columns retain RNA but not protein-crosslinked RNA under standard conditions, we subjected XRNAX extracts to limited tryptic digestion resulting in efficient retention of RNA, which was now crosslinked to protein fragments. This was evidenced upon full proteolysis, resulting in the identification of a much larger number of peptides from crosslinked cells (Figure 1F). The majority of all peptides had an intensity ratio >1000 of crosslinked over non-crosslinked and in the following we refer to these as ‘super-enriched peptides’.

In a second validation we tested if silica enrichment depended on RNA. Therefore, we degraded RNA in an XRNAX extract from light SILAC cells with NaOH/Mg^2+^. After pH-neutralization we combined it with an untreated XRNAX extract from heavy SILAC cells and subjected the mixture to the workflow described above. Consequently, we identified many thousands of super-enriched peptides originating from the sample with intact RNA, showing that their enrichment via silica columns strongly depended on the integrity of RNA (Figure S1D).

In a third validation we established that super-enriched peptides could be used to quantify changes in protein-RNA interactions. To this end, we mixed decreasing amounts of UV-crosslinked MCF7 cells with non-crosslinked cells both grown with heavy SILAC label, and combined this with a fixed amount of UV-crosslinked cells of a light SILAC label. Thereby, these samples contained discrete ratios of RNA-crosslinked over non-crosslinked proteins ranging from 1:4 to 1:256 (Figure S1E). Indeed, after processing samples via XRNAX and silica enrichment, most peptides were accurately quantified according to their dilution over two orders of magnitude (Figure S1E). A comparison to our earlier data (Figure 1F) revealed that peptides that showed these discrete fold changes were exclusively found among the super-enriched peptides. Conversely, peptides that remained constant irrespective of the mixing ratio (Figure S1E) were found among the less-enriched peptides and disregarded for further use. This demonstrated that XRNAX in combination with silica enrichment could be used to quantify RNA-binding of proteins between conditions.

In numerous validation experiments using a fluorescence-based system or RNA radioloabeling through polynucleotide kinase (PNK), respectively, we(Castello et al., 2012) and others(Baltz et al., 2012) have established that protein UV-crosslinked to RNA represents direct evidence for protein-RNA interactions and can be used to screen for novel RNA-binding proteins or interaction sites. Our experiments indicated that XRNAX extracts were highly enriched in RNA-binding proteins, which were UV-crosslinked to RNA. Furthermore, conventional silica chromatography could be used to purify some fragments of UV-crosslinked proteins to the point, where there was effectively no background detectable. Consequently, these fragments, represented by super-enriched peptides, could not only be used to classify RNA-binding proteins in a very conservative way but also to quantify RNA-binding differentially.

Having established XRNAX as an initial extraction method for accessing protein-crosslinked RNA we utilized it in three designated applications outlined in Figure 2A, all of which use 254 nm UV-crosslinking to characterize and validate the composition of the protein-RNA interactome.

**Figure 2:**
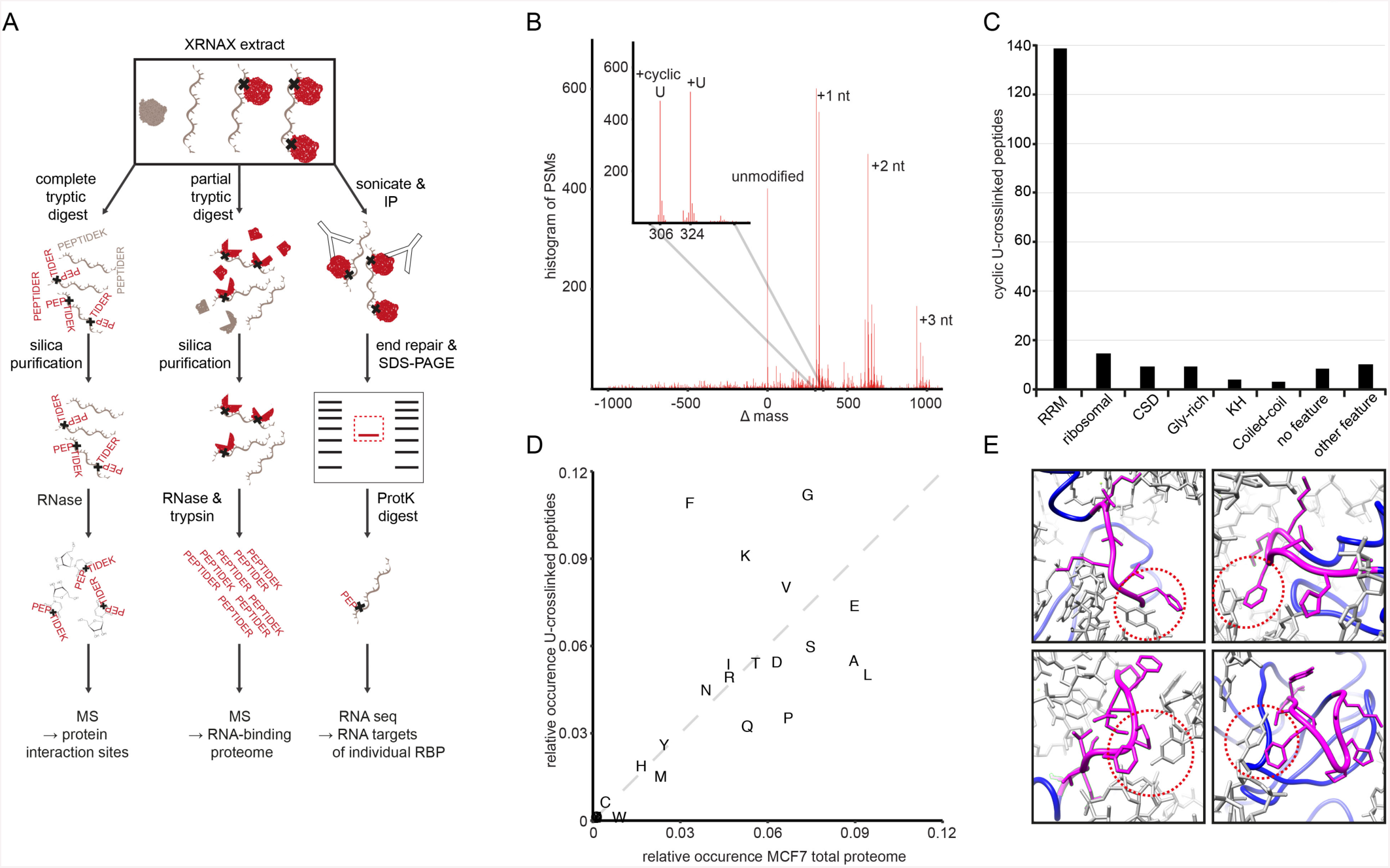
Identification of RNA interaction sites and domains. (A) Proteomic and transcriptomic applications downstream of XRNAX for the (B)Identification of crosslinking sites by MS (left), the characterization of RNA-bound proteomes (middle), and identification of protein-bound RNA by CLIP-Seq. Identification of adduct masses on XRNAX-purified peptides by MSFragger. (C) Uniprot feature-annotation of cyclic U-crosslinked peptides. Bar graph displays number of cyclic-U crosslinked peptides that were found in stretches of their cognate protein with indicated feature. RRM: RNA recognition motif; ribosomal: host protein is constituent of the ribosome; KH: K homology domain; CSD: cold shock domain; Gly-rich: glycine-rich amino acid sequence; no feature: no feature deposited in Uniprot; other feature: feature other than the ones mentioned in the other categories. (D) Comparison of amino acid frequencies in peptides crosslinked to cyclic U and peptides found in the MCF7 total proteome. (E) Protein-RNA interactions in crystal structure of the human ribosome for four exemplary proteins. The amino-acid backbone is indicated in blue, peptides found crosslinked to cyclic U are indicated in magenta and RNA is indicated in grey. Red circles highlight phenylalanine in the vicinity of a uracil base. Upper left: GFVKVVK in RPL5, upper right: IHGVGFKK in RPL31, lower left: MKFNPFVTSDR in RPL26L1, lower right: VAYVSFGPHAGK in RPL14.

### Direct Detection of Nucleotide-Crosslinked Peptides from XRNAX Extracts

First, we used MS to identify RNA-crosslinked peptides and RNA interaction sites (Figure 2A, left). Direct identification of RNA-modified peptides has proven challenging as UV-induced RNA-protein crosslinking happens only sparsely, and hybrids are not easily purified from non-crosslinked peptides and ribonucleotides (Darnell, 2010; Kramer et al., 2014). Moreover, any of the four ribonucleotides may crosslink to any of the 20 amino acids, thus tremendously inflating the search space when interpreting MS data. To circumvent this, previous methods like RBR-ID (He et al., 2016) used the absence of peptide-RNA hybrids in MS data to infer protein-RNA interaction sites, whereas RBDmap identified peptides in the direct neighborhood of those hybrids (Castello et al., 2016). Alternatively, by making *a priori* assumptions for the chemical adduct between ribonucleotide and amino acid, Kramer et al. successfully identified 60 nucleotide-crosslinked peptides from 35 human proteins (Kramer et al., 2014). Here, we aimed to take an unbiased approach for the direct localization of RNA-protein crosslinking sites, taking advantage of XRNAX to enrich for RNA-crosslinked proteins. Therefore we digested the XRNAX extract from MCF7 cells with an excess of trypsin, purified RNA-bound peptides via silica columns, digested RNA and analyzed resulting peptides by MS (Figure 2A). Importantly, for the unbiased identification of any remaining nucleotide adducts, we used the recently reported mass-tolerant search engine MSFragger (Kong et al., 2017). This identified 324 and 306 Dalton as the most dominant mass shifts (Figure 2B), corresponding to peptide modifications by uridine monophosphate (U) and cyclic uridine monophosphate (cyclic U), respectively. Mononucleotide-peptide adducts of U were in accordance with the earlier study by Kramer et al., however, in addition we identified oligonucleotide-adducts consisting of permutations of di- and trinucleotide sequences carrying at least one U (Figure 2B). Based on this we suggest that U acts as the main crosslinking base, although we cannot exclude that crosslinking with other ribonucleotides escaped MS/MS detection. Collectively, this led to the detection of 197 cyclic U-crosslinked peptides from 93 proteins (Table S1), including the ones that were known before (Kramer et al., 2014). These findings provided direct evidence that XRNAX extracts contained RNA-crosslinked proteins, while showing that MSFragger identified protein-RNA interfaces in an unbiased way.

### Nucleotide-Crosslinked Peptides Locate Known and Novel Protein-RNA Interfaces and Identify DUF2373 as a Potential RNA-Binding Domain

Of the 93 proteins that were directly identified by nucleotide-crosslinked peptides, 90 % were annotated as RNA-binding (Table S1). In addition, more than 85 % of nucleotide-crosslinked peptides mapped to *bona fide* RNA-binding domains, confirming that these peptides represented protein-RNA interfaces and that the identification of crosslinks validated interaction sites. Most of the peptides localized in RNA-recognition motif (RRM), K-homology (KH) or cold-shock domains (CSD) (Figure 2C, Table S1). In addition we frequently identified glycine-rich regions, in accordance with earlier RBDmap and RBR-ID studies showing that low-complexity regions are abundantly involved in RNA-binding (Castello et al., 2016; He et al., 2016). Proteins identified with the most crosslinked peptides were HNRNPA2B1 (13), NCL (10) and HNRNPAB (8), all of which located to RRMs or glycine-rich regions (Figure S2A).

We identified the SRA RNA-interactors SLIRP and SPEN (also known as SHARP, MINT) and located the RNA interaction site of SPEN to its RRM3, which reportedly is essential in SRA RNA-binding (Arieti et al., 2014). RBDmap had located the interaction site of SPEN with polyadenylated RNA to its RRM1 (Castello et al., 2016), suggesting alternative binding modes depending on the RNA substrate. In another instance we identified a crosslinked peptide in the SLED domain of the polycomb group member SCMH1 (Table S1). The SLED domain has been shown to allow for sequence-specific binding of double-stranded DNA (Bezsonova, 2014), which according to our finding may be modified through interaction with RNA. We encountered several cases where cyclic U-crosslinked peptides located to non-canonical RNA-binding domains, pointing out interesting structural or functional properties. For example, we located the RNA interface of the ribosome biogenesis factor LTV1 to a C-terminal coiled-coil region (Table S1). Although no RNA-binding domain is known for LTV1, cryo-electron microscopy studies indicate this region in the interaction with ribosomal RNA during pre-40S ribosome assembly (Larburu et al., 2016). In yet another instance we located a cyclic U-crosslinked peptide in the AAA+ ATPase domain of HNRNPU (also known as SAF-A) (Table S1). Only recently a study reported the importance of this domain in HNRNPU’s role in chromosome organization, and *in vitro* experiments with the isolated domain indicated that its ATP hydrolyzing activity is indeed stimulated in the presence of RNA (Nozawa et al., 2017). Our findings implied that the C-terminal region of the AAA+ ATPase domain interacted with RNA, suggesting that this stimulation might occur through RNA directly.

Interestingly, we identified a nucleotide-crosslinked peptide in the domain of unknown function DUF2373 of the uncharacterized protein C7orf50 (Table S1). We previously had identified C7orf50 as an RNA-binding protein in HeLa cells (Castello et al., 2012), however, it did not carry any known RNA-binding domains, nor did it appear in RBDmap or RBR-ID studies (Castello et al., 2016; He et al., 2016). Analysis in STRING showed that C7orf50 interacts with DKC1, TRUB1 and FTSJ3 (Figure S2C) relating it to ribosome biogenesis and pseudouridylation, which was supported by nucleolar localization reported in the Human Protein Atlas (Bailey et al., 2009) (Figure S2D). We propose renaming DUF2373 as the WKF domain after the three most conserved amino acids in the sequence alignment (Figure 2B). This naming scheme has been adopted by the Pfam database. The WKF domain is found widely in animal and fungal species suggesting that the protein has an important yet unknown function.

In summary, these data significantly expanded the number of proteins with direct MS evidence for RNA interaction. Furthermore, the identified protein-RNA interfaces suggested novel RNA-binding modes, gave structural insights, hypothesized mechanisms for the modulation of enzymatic activity and provided evidence for a novel RNA-binding domain.

### Uracil-Phenylalanine Interactions Are Important During UV-Crosslinking

Analysis of amino-acid frequencies in peptides crosslinked to cyclic U showed a clear enrichment for phenylalanine, lysine and glycine compared to the total MCF7 proteome (Figure 2D). We searched our cyclic U-crosslinked peptides with the motif search-engine MEME and identified the conserved RNP1 or RNP2 motif in 71 % of cases. Both motifs are rich in phenylalanine and are known to be critical for RNA recognition of the RRM (Elliott and Ladomery, 2015), suggesting the importance of phenylalanine in those interfaces during UV-crosslinking. Indeed, a recent MS study identified phenylalanine in the RRM of the splicing factor PTBP1 to crosslink with uracil in RNA (Dorn et al., 2017). Moreover, Kramer et al. had discussed stacking interactions between the crosslinking base and aromatic side chains from the crystal structure of mRNA-binding proteins. We could now explore this more broadly, benefiting from the ability of XRNAX to recover other RNA types such as rRNA. In the crystal structure of the 80S human ribosome, we could locate cyclic U-crosslinked peptides from seven ribosomal proteins (Table S1, Figure 2E). Notably, for four out of six peptides that contained phenylalanine, we found phenylalanine in close proximity to a uracil base in the RNA sequence. In conclusion, our data suggest that phenylalanine-uracil adducts may be a preferred form of UV-induced protein-RNA crosslink.

### An Integrated Draft of the Human RNA-Binding Proteome From Three Cell Lines

To explore the complete RNA-bound proteome, we applied our initial enrichment scheme (XRNAX and partial digestion followed by silica enrichment, Figure 2A, center) to three commonly used human cells lines, MCF7, HeLa and HEK293, which had been subject of poly(A)-interactome capture studies before (Baltz et al., 2012; Castello et al., 2012; Milek et al., 2017). To classify proteins identified as RNA-binding we only considered proteins that were detected with at least two super-enriched peptides, i.e. with a SILAC-ratio >1000, to reflect that they were exclusively found in crosslinked cells (Figure 1F), in a reproducible manner (Figure 3A). This resulted in high-confidence RNA-binding proteomes of 1207 proteins for MCF7, 1239 proteins for HeLa and 1357 proteins for HEK293 cells, of which 858 were shared by all three cell lines (Figure 3B, Table S2).

**Figure 3:**
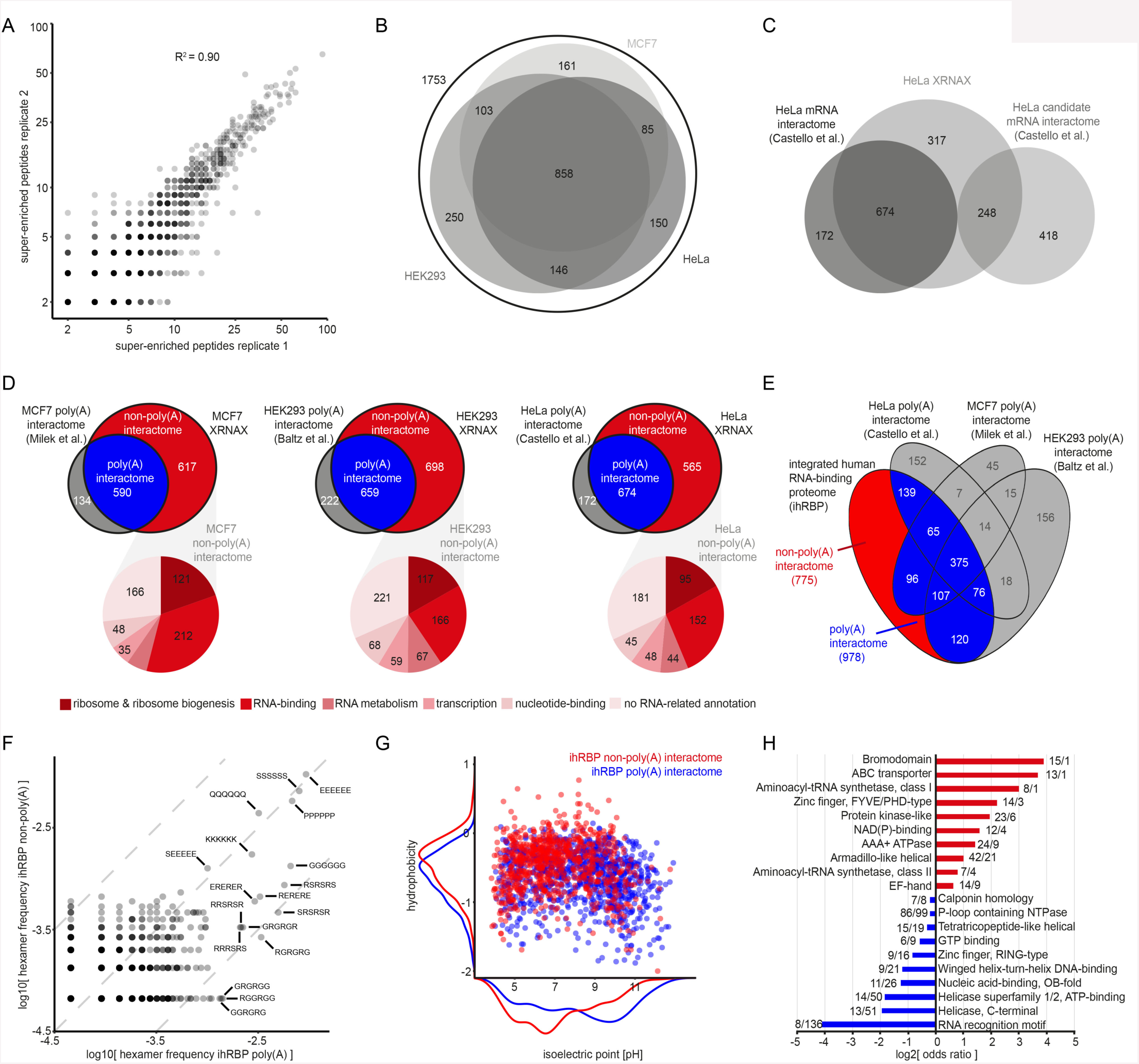
The integrated human RNA-binding proteome derived from 3 cell lines. (A) Scatter plot showing reproducibility of protein identifications by super-enriched peptides. Each point represents one protein and indicates how many super-enriched peptides were found per replicate. Note logarithmic scaling of axis. (B) Venn diagram for XRNAX-derived RNA-binding proteomes. Numbers indicate the proteins identified in each of the three cell lines MCF7, HeLa and HEK293. (C) Venn diagram comparing the XRNAX-derived HeLa RNA-binding proteome to the published poly(A) RNA interactome by Castello et al. (2012). (D) Composition of XRNAX-derived RNA-binding proteomes. Top: Venn diagrams comparing XRNAX-derived RNA-binding proteomes to published poly(A) interactomes. Non-poly(A) interactomes (red) were derived by subtraction of poly(A) interactomes (blue). Bottom: Pie chart for the functional annotation of proteins in the non-poly(A) interactome into five RNA-related functional categories. (E) Venn diagram comparing the integrated human RNA-binding proteome (ihRBP) to published poly(A) interactomes. A non-poly(A) RNA interactome (red) was derived by subtraction of the combined known poly(A) interactomes (blue). (F) Scatter plot comparing hexamer frequencies in poly(A) and non-poly(A) interactomes. All possible hexameric permutations of the amino acids G, S, N, Q, P, E, K and R were counted in proteins of each group and normalized to the total number of counts. Dashed lines indicate fold-changes of 1 and 10. (G) Scatter plot comparing isoelectric points and hydrophobicity of proteins in poly(A) and non-poly(A) interactomes. Density plots outside the axes illustrate the distribution for each feature. (H) Odds ratios of interpro domain occurrences in poly(A) and non-poly(A) interactomes. The ten most frequent domains in either group are compared.

The XRNAX-derived HeLa RNA-binding proteome rediscovered 647, i.e. 80 %, of proteins found in our previous poly(A)-interactome capture (Castello et al., 2012) and added almost 600 more (Figure 3C). Although we applied a more stringent enrichment cut-off for calling RNA-binding proteins (presence-absence instead of fold-change relative to a non-crosslinked control), we were able to confirm some of the proteins that were earlier reported as ‘candidate mRNA-binder’ (Figure 3C) because they had failed to reach statistical significance (Castello et al., 2012). Similarly, XRNAX-derived RNA-binding proteomes in MCF7 and HEK293 cells showed large (>75%) overlap with their published poly(A) interactomes (Figure 3D, top). Furthermore, in each cell line we identified more than 600 additional proteins, which here we refer to as the non-poly(A) interactome.

We were interested to see how much of the non-poly(A) interactomes could be explained by current GO annotations. For this purpose we designated five RNA-related categories summarizing several related GO terms, namely ribosome & ribosome biogenesis, RNA-binding, RNA metabolism, transcription and nucleotide binding. Since many RNA-binding proteins were represented in several of those categories, we applied them hierarchically and removed proteins from our list as soon as they fit into one category (Figure 3D, bottom). In this way, more than 70% of proteins in the non-poly(A) interactomes could be assigned. This means that among the entire XRNAX-derived poly(A) and non-poly(A) interactomes, more than 80% of all proteins had a prior annotation related to the interaction with RNA or ribonucleotides, testifying to the specificity of XRNAX.

Combining our findings from all three cell lines resulted in a collection of 1753 proteins that we named the integrated human RNA-binding proteome or ihRBP, containing 978 proteins (70 %) of previous poly(A) interactomes, and 775 proteins constituting the novel non-poly(A) interactome (Figure 3E). Comparison of the ihRBP to the census of RNA-binding proteins from Gerstberger et al., who had catalogued RNA-binding proteins by combining computational analyses with manual curation (Gerstberger et al., 2014) showed 55 % overlap (Figure S3A).

This illustrated an observation made in earlier studies that computational prediction of RNA-binding is often incomplete, because RNA-binding can be mitigated by features outside classical domains such as intrinsically disordered regions (IDRs) (Beckmann et al., 2015; Castello et al., 2012, 2016).

GO analysis of the ihRBP revealed strong enrichment for RNA-related terms in the poly(A) interactome (Figure S3B), using deep total proteome data as a background (Geiger et al., 2012). A similar analysis was problematic for the non-poly(A) interactome because of the high prevalence of RNA-binding proteins in the total proteome of the three cell lines, annotated as such in reference to the two initial poly(A) interactome studies by Baltz et al. and Castello et al.. Since for this particular analysis we were exclusively interested in non-poly(A) interactors, we removed the published poly(A) interactome from the background control. Consequently, we found strong enrichment for RNA-related terms in the non-poly(A) interactome as well (Figure S3B). Interestingly, terms relating to non-coding RNA such as rRNA, 7S RNA, tRNA, snRNA or snoRNA and the ribosome were especially enriched. Moreover, some terms such as ‘aminoacyl-tRNA ligase activity’ or ‘ribonuclease activity’ were found only enriched in the non-poly(A) part of our interactomes. Indeed, the non-poly(A) interactome of the ihRBP contained numerous proteins known to interact with non-coding RNAs, such as ribosomal proteins, numerous ribosome biogenesis factors (e.g. LTV1, MDN1, RIOK1-3, RIOX2), 17 out of 23 aminoacyl tRNA synthetases, translation initiation factors (e.g. ABCE1, EIF2S3, EIF3B, EIF3J), RNA exosome components (e.g. SKIV2L, EXOSC2, EXOSC3, DIS3), splicing factors (e.g. SF3B3, LSM8, ESRP1 or CWC22), proteins involved in micro RNA biogenesis (e.g. DICER1, TSN and TARBP2), many transcription-associated proteins e.g. known to interact with 7SK RNA (e.g. HEXIM1, CCNT1, CDK9), and interactors of telomerase RNA (NOP10, PINX1).

Thus, XRNAX in combination with silica enrichment allowed us to identify highly reproducible, high-confidence RNA-binding proteomes. These included the large majority of the known poly(A)-interactomes and added more than 600 proteins, many of which are known interactors of ncRNA.

### Sequence-Encoded Information Distinguishes Poly(A) from Non-Poly(A) RNA Interactors

We next asked if any sequence-encoded properties could be identified that distinguished poly(A) from non-poly(A)-binding proteins in the ihRBP. While amino acid frequencies were virtually identical between the two groups (Figure S3C), distinct differences became apparent when comparing di- and tripeptide frequencies within proteins (Wilcox ranksum test p=0.02). Interestingly, proteins of the poly(A) interactome showed the strongest enrichment for tripeptides carrying combinations of amino acids known to contribute to IDRs (Figure S3C), i.e. glycine (G), serine (S), asparagine (N), glutamine (Q), proline (P), glutamic acid (E), lysine (K), and arginine (R) (Brangwynne et al., 2015). To get a more detailed picture, we limited the analysis to these eight IDR amino acids and compared hexamer frequencies between the two groups (Figure 3F, Figure S3D). This specifically highlighted hexapeptides consisting of alternating R and E, R and S, R and G, as well as hexa(G) in the poly(A) interactome. Increased frequency of low-complexity motifs in poly(A)-binding proteins has been reported before (Calabretta and Richard, 2015; Castello et al., 2016). However, our analysis revealed for RNA-binding proteins in general that, depending on the RNA biotype they bind to (i.e. poly(A) vs. non-poly(A)), they contained specific low-complexity motifs more frequently. Other motifs, such as poly(E), poly(S), poly(P), poly(Q) or poly(K) stretches occurred with very similar frequencies in both groups. Interestingly, proteins carrying those repeats were often annotated with similar molecular processes, for example the GO term ‘chromatin organization’ was enriched among proteins carrying a poly(E) stretch.

When comparing hydrophobicity, isoelectric point, molecular weight and charge state at pH=7 between the two groups, we found that poly(A) binders were generally more hydrophobic and had a more alkaline isoelectric point (pI), whereas non-poly(A) binders were larger and often carried more negative charges (Figure S3E). While the differences in physicochemical features were highly significant, Figure 3G illustrates that the variance was high, so that no single feature could be used for effective classification between the two groups. This suggested that IDRs and the superposition of physicochemical characteristics could guide the assembly of poly(A) or non-poly(A) ribonucleoprotein particles, respectively.

### Analysis of Protein Domains Highlights Chromatin Remodelers as Novel RNA-Binders

A comparison of domain structure revealed notable commonalities and differences between the poly(A) and the non-poly(A) interacting proteomes (Figure 3H). P-loop-containing nucleoside triphosphate hydrolases represented the largest, albeit equally distributed domain superfamily. RRM-containing proteins were almost exclusively found in the poly(A) interactome, which was also enriched for RING-type zinc finger domains. In contrast, zinc finger domains of the FYVE/PHD-type were much more frequent among non-poly(A) binders. As expected, we found both classes of aminoacyl-tRNA synthetase domains strongly enriched in the non-poly(A) interactome, which also contained an overrepresentation of the ABC-transporter-like and the AAA+ ATPase domains. Surprisingly, the domain with the strongest enrichment in the non-poly(A) interactome was the bromodomain, which only recently had been recognized among RNA-binders (He et al., 2016). The absence of these proteins in poly(A)-interactome studies and their presence in RBR-ID and XRNAX, which both target RNAs globally, suggested that bromodomain proteins do in fact interact with non-polyadenylated ncRNA.

These observations prompted us to look more systematically for chromatin-modifying proteins in the non-poly(A) interactome using the STRING interaction database. For instance, we identified TP53 and the DNA-damage regulators TP53BP1, BAX, FANCI, RPA1, DDB1, RIF1, MDC1, as well as BRCA1. TP53BP1 (Francia et al., 2012) and BRCA1 (Ganesan et al., 2002) has been reported to interact with non-coding RNA, whereas RNA-binding by TP53 has been controversial (Riley and Maher, 2007). Notably, we detected 25 super-enriched peptides for TP53BP1, ranking it among the most confidently classified non-poly(A) interactors.

Other clusters contained proteins involved in chromosome segregation (such as BUB3, AHCTF1, CKAP5, PDS5A, KIF2C, KIF11 and CENPF) and around the condensin complex (such as SMC2, SMC4, NCAPD2, NCAPD3, NCAPDG and NCAPG2), none of which had been reported before to bind RNA. These findings suggested that also proteins involved in cytokinesis bind RNA, which could be a way of preserving chromosomal integrity outside the cell cycle interphase (Nozawa et al., 2017).

Finally, a cluster of proteins around POLR1A contained CHD1 (a member of the SAGA complex), CHD8, CDK9, ATRX (member of the ATRX:DAXX complex), TOP2B, and the RNA polymerase I transcription termination factor TTF1. Except for POLR1A, we could not find previous reports for any of the proteins in this cluster to interact with RNA. As for bromodomain proteins, proteins in this cluster are closely involved in transcription and transcription regulation, again suggesting a direct link of non-poly(A) interaction and gene expression.

As non-coding RNAs emerge as important regulators in genome regulation (for review see (Long et al., 2017; Wang and Chang, 2011) future studies may investigate if the function of chromatin components identified here could be guided by RNA.

### Arsenite Induces Translational Arrest and Autophagic Remodeling of the Translation Machinery

We next applied XRNAX to investigate changes in the protein-RNA interactome in arsenite-treated MCF7 cells. Arsenite stress has been thoroughly studied in the context of protein-RNA interactions during translational arrest and the formation protein-RNA complexes known as stress granules (for review see (Anderson and Kedersha, 2009; Buchan and Parker, 2009)). While the composition of arsenite-induced stress granules (Jain et al., 2016; Khong et al., 2017) and the effect of arsenite-induced translational arrest on the transcriptome (Andreev et al., 2015) are well understood, very little is known about its effect on the proteome. In order to get a comprehensive understanding of the cellular responses to arsenite treatment, we investigated its effect on the RNA-bound proteome and integrated this with effects on translation and the total proteome. To this end, we first performed a time course experiment to select the shortest possible timescale to capture immediate consequences of arsenite stress, and monitored protein biosynthesis to confirm arsenite-induced translational arrest. Specifically, we used SILAC and azidohomoalanine (AHA) labeling to isolate nascent proteins by click-chemistry and quantify them by MS as a direct measure for translation (Eichelbaum and Krijgsveld, 2014; Eichelbaum et al., 2012).

Indeed, translation was heavily decreased in arsenite-treated cells after 60 minutes, down to 23% (median) compared to the untreated control (Figure 4A). Moreover, this occurred at a remarkably rapid pace, already apparent after ten minutes and gradually reaching its maximum at 30 minutes.

**Figure 4:**
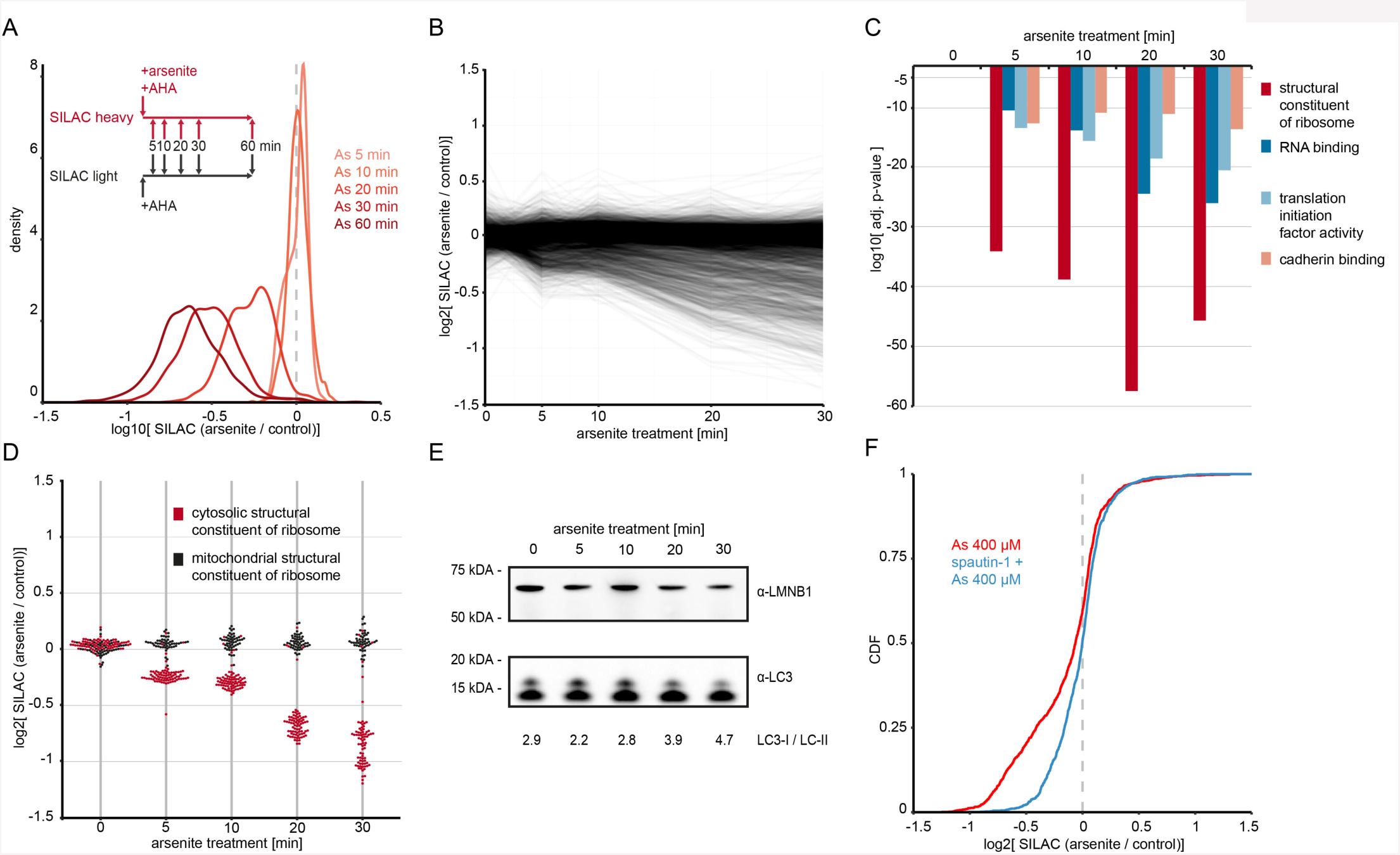
Impact of arsenite stress on the proteome. (A) Density plot showing fold-changes of nascent proteins produced under normal conditions or arsenite stress. MCF7 cells were treated with arsenite and simultaneously nascent protein labeled by incorporation of AHA. Cells were harvested at the indicated times, and levels of newly synthesized proteins were compared to a non-treated control. Displayed are means of duplicate experiments with label-swap. (B) Time course of total proteome changes during arsenite stress in MCF7 cells. (C) GO-analysis of proteins that change in expression during arsenite stress. Shown are the top-4 GO terms with highest significances after 30 minutes. (D) Dotplot displaying changes in total proteome upon arsenite stress for proteins under the GO term ‘structural constituent of ribosome’. Each dot represents one protein. Values are means of duplicate experiments with label swap filtered for a variance of 15 % or smaller. (E) Western blot against LC3 to monitor autophagic flux in MCF7 cells over 30 minutes of arsenite treatment. LMNB1 was used as a loading control. Numbers under images quantify the relative intensity of LC3-I and LC3-II bands. (F) Cumulative distribution of changes in total proteome of MCF7 cells upon arsenite treatment, with and without inhibition of autophagy by spautin-1. Cells were treated for 24 hours with 10 µM spautin-1 (blue) or left untreated (red) before heavy SILAC labelled cells were treated with arsenite for 30 minutes, compared to light SILAC labelled cells without arsenite treatment (see also Figure S4G).

Next, we analysed changes in overall protein expression levels over five time points within the first 30 minutes of arsenite treatment. This revealed that most of the proteins were unaffected, however, part of the proteome showed a distinct and gradual decrease in protein abundance (Figure 4B, Table S3). We performed GO analysis for downregulated proteins and found profound enrichment for translation-related terms emerging already after five minutes (Figure 4C). Closer examination of ribosomal proteins, as the most prominently affected group, revealed the specific decrease of cytosolic but not mitochondrial ribosomes (Figure 4D). This was concomitant with the collective downregulation of eukaryotic translation initiation factors (EIFs) with similar kinetics, effectively reducing their expression level to 50% within 30 minutes (Figure S4A). In addition, other RNA-binding proteins decreased in abundance, most of them with functionalities in protein biosynthesis (Figure S4B, Table S3).

Observing the highly selective degradation of a narrow portion of the proteome, we considered that this may be regulated via autophagy. Indeed, arsenite-induced protein degradation was markedly decreased upon pre-treatment with the autophagy inhibitor spautin-1 (Figure 4F), while this was not the case after pre-treatment with the proteasome inhibitor bortezomib (Figure S4C). Western blotting against the autophagy marker LC3 confirmed increased autophagic flux over the course of arsenite stress (Figure 4E).

In conclusion, we uncovered that arsenite induced a severe reduction in translation within 30 minutes. This was concomitant with profound, rapid, and highly specific remodeling of the proteome, invoking autophagy to selectively degrade proteins operating in the translation apparatus.

### Exploring The Dynamics of Protein-RNA Interactions During Arsenite Stress

We next investigated how arsenite stress induced changes in the RNA-bound proteome, and compared this to global proteome changes identified above. To do so, we challenged MCF7 cells of one SILAC label with arsenite and compared this to untreated cells of the complementary label. Using XRNAX followed by silica enrichment and MS, we recorded data for five time points in a 30 minutes window. After filtering for the super-enriched peptides identified previously, we quantified the association of 765 proteins with RNA over all time points (Figure 5A and 5B, Table S4). While most proteins did not change their RNA-binding under arsenite stress (90 % quantiles after 30 minutes were within 0.77 - 1.21 fold change), several proteins showed significantly decreased association with RNA, whereas the only protein increasing more than twofold was TP53BP1 (Figure 5B). The kinetic profile of these proteins showed a steady incline or decline, with the exception of the RNA exosome component EXOSC2, which already showed increased RNA-binding after five minutes of arsenite stress, and then stayed constant.

**Figure 5:**
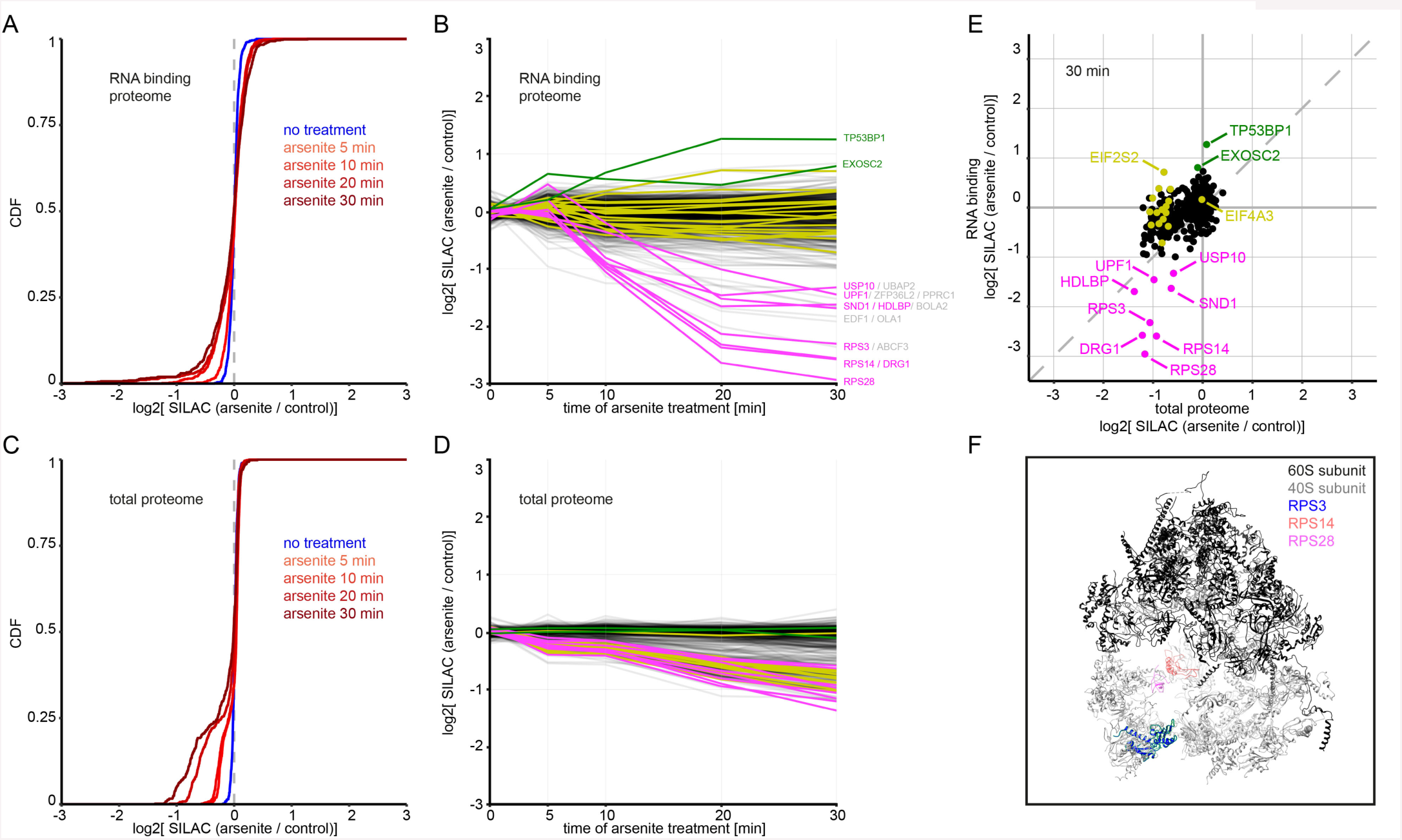
Changes in the total and RNA-interacting proteome induced by arsenite-mediated translational arrest. (A) Cumulative changes in the RNA-binding proteome, assessed by XRNAX in MCF7 cells that were treated with arsenite for indicated times. (B) As in panel A, showing temporal data for individual proteins. Each line represents one protein. Values displayed are means of duplicate experiments with label swap filtered for a variance of 15 % or smaller. (C) Cumulative distribution of changes in the total MCF7 proteome upon arsenite stress for proteins displayed in panel A. (D) Timeline of changes in the total proteome upon arsenite stress for proteins displayed in panel B. (E) Scatter plots comparing changes in the total proteome to changes in RNA-binding after 30 minutes of arsenite stress. For other time points see Figure S5. Color coding in panels B, D and E refers only to proteins quantified in both the RNA-binding and total proteome. The proteins TP53BP1 and EXOSC2 are displayed in green, proteins with >50 % change in RNA-binding after 30 minutes are displayed in magenta, and eukaryotic translation initiation factors (EIFs) are displayed in yellow. (F) Crystal structure of the human ribosome highlighting the location of RPS28, RPS14 and RPS3. For visibility nucleic acids are not displayed.

In order to control for total protein abundances, we intersected this dataset with our data for the total proteomes, which resulted in 619 proteins that were quantified over all time points (Figure 5C-E, see also Figure S5A). Figure 5C illustrates that about 25 % of the quantified RNA-binding proteome was affected by the autophagic degradation process, which we had characterized earlier (Figure 4B). Consequently, a direct comparison of RNA-binding to protein expression revealed several remarkable patterns. First, the increase in RNA-binding of the proteins TP53BP1 and EXOSC2, could be entirely attributed to their association with RNA, because their absolute abundance stayed constant over time (Figure 5E). Second, the proteins showing the strongest reduction in RNA-binding also decreased in protein abundance (Fig 5E, magenta). However, this decrease was small in comparison to their decreased RNA binding, primarily indicating a release from RNA. Interestingly, this included the ribosomal proteins RPS28, RPS14 and RPS3, which are all positioned in the cleft of the 80S-ribosome that directly interacts with mRNA to channel it through the two ribosomal subunits (Figure 5F). Thus, the measured decrease in the interaction of core ribosomal components with RNA recapitulated the disassembly of polysomes during the progression of translational arrest.

A third striking observation was the 40-50 % decrease in protein abundance of nearly all detected eukaryotic translation initiation factors (EIFs), without a significant change in RNA binding (Figure 5E, yellow, see also Figure S5B). A similar observation could be made for cytosolic ribosomal proteins other than the ones mentioned above (Table S4), suggesting that RNA-binding might protect from protein degradation.

Two exceptions here were EIF4A3 and EIF2S2. EIF4A3 is one of the RNA-binding components of the exon junction complex and thereby only tangentially involved in translation initiation (Shibuya et al., 2004), which might explain why it did not share the same behaviour as the other EIFs. EIF2S2, also known as eIF2-α and core component of the EIF2 complex involved in 43S preinitiation complex formation (for review see (Jackson et al., 2010)), increased RNA-binding steadily over all time points, although its protein abundance decreased significantly (Figure S5B). In fact, by normalizing RNA-binding with protein abundance its effective 2.8-fold increase in RNA-binding was more pronounced than for any other protein in our data. This might indicate that 43S preinitiation complexes assembled on RNA, while translational arrest occurred downstream of this process.

Lastly, another protein with effectively reduced RNA-binding was USP10, which reportedly has a key role in stress granule formation (Anderson and Kedersha, 2009). These observations were not affected by changes in the integrity or amount of RNA, since total RNA from arsenite-treated MCF7 cells was neither subject to degradation (Figure S4D and S4E) nor altered turnover (Figure S4F), in line with previous transcriptomic data(Andreev et al., 2015).

In summary, our data illustrated that quantification of RNA-binding using XRNAX recapitulated RNA-ribosome dissociation known to occur during translational arrest, and added molecular detail on the association of EIFs with RNA. In addition it identified RNA-binding proteins previously not known to be involved in the cellular response to arsenite, suggesting novel players and mechanisms that future studies may address in more detail.

### Linking XRNAX with CLIP-Seq to Identify Protein-Binding RNAs

Having successfully employed XRNAX for the proteomic analysis of protein-RNA interactions, we next aimed to combine it with CLIP-seq to validate some of these data by identifying RNAs that interact with proteins identified in the course of this study. Conceptually, XRNAX as a sample preparation step prior to CLIP-Seq is advantageous for a number of reasons: i) Contaminants like DNA, which could physically obstruct immunoprecipitation or mask target protein in chromatin complexes, are eliminated. ii) Sample volumes are reduced from milliliters to microliters, thereby allowing for higher antibody concentrations. iii) RNA fragmentation can be supplemented by high-intensity sonication, thereby circumventing cumbersome optimization and potential biases of RNase treatment (Haberman et al., 2017).

We selected Lamin B1 (LMNB1) as a CLIP target, to validate it as a novel RNA binder, which we had identified among the proteins with the highest number of super-enriched peptides in the non-poly(A) interactome of MCF7 cells (Table S2). We fragmented RNA in an XRNAX extract using ultrasonication, and immunoprecipitated LMNB1 using a variation of the eCLIP protocol ((Van Nostrand et al., 2016), Fig. S6A, for details see methods). RNA sequencing identified various ncRNAs that were significantly enriched over the size-matched input control, primarily snoRNAs but also other small nuclear RNAs (Figure 6A). This was in agreement with previous reports showing that Lamin B, with a canonical function in the nuclear lamina, is also structural component in nucleoli where its presence is required to maintain nucleolar integrity during ribosome biogenesis (Martin et al., 2009). Whether the interaction between Lamin B1 and snoRNAs fulfills a function may be the topic of future studies.

**Figure 6:**
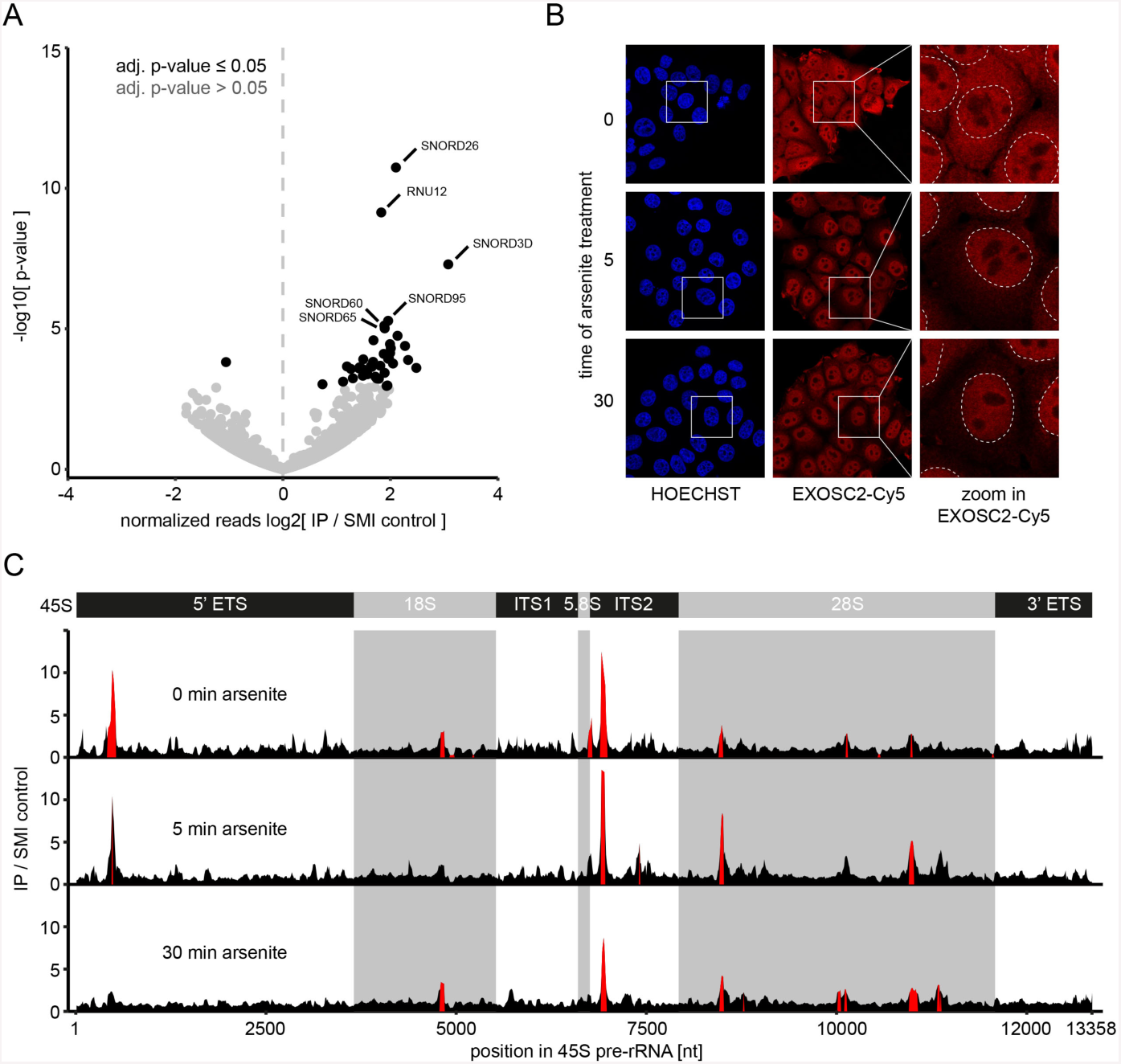
Combination of XRNAX with CLIP-seq to identify RNA bound to novel RNA-binding proteins. (A) Volcano plot showing the enrichment of non-coding nuclear transcripts in XRNAX CLIP-seq for LMNB1 in MCF7 cells. (B) Immunoflourescence detection of EXOSC2 with confocal microscopy during translational arrest. White squares in the left and middle images mark the magnified areas displayed on the right. White, dashed lines in images on the right indicate the outline of nuclei. (C) Gene track for the 45S pre-RNA displaying coverage of XRNAX-CLIP-seq for EXOSC2, normalized to the SMI control. IP coverage enriched over the SMI control with an adj. p-value < 0.001 is highlighted in red (for details see methods). Top scheme indicates location of transcripts that get processed from the 45S premature transcript. Grey shading refer to the mature 18S, 5.8S and 28S transcripts. 5’ ETS: 5’ external transcribed spacer, 3’ ETS: 3’ external transcribed spacer, ITS1: internal transcribed spacer 1, ITS2: internal transcribed spacer 2.

Methodologically, these experiments showed that XRNAX extracts can serve as direct input for CLIP-seq experiments, and thereby demonstrated that XRNAX can be applied both for the discovery and validation of novel RNA-binding proteins by MS and RNA-sequencing, respectively, from the same sample.

### Differential RNA-Binding of EXOSC2 Upon Arsenite Stress

Our differential quantification of RNA-binding had shown that EXOSC2 (also known as RRP4) increased its interaction with RNA already after five minutes of arsenite stress (Figure 5B, Table S4). Since this temporal behaviour stood out among all other proteins, we decided to investigate further. EXOSC2 is a component of the non-catalytic ‘lid’ of the exosome, which is distinct from the catalytic core that degrades RNA (Zinder and Lima, 2017). In fact, we also detected the core protein EXOSC10, which exhibited a similar yet less pronounced RNA-binding kinetic than EXOSC2 (Table S4). We performed immunohistochemistry and confocal microscopy to locate EXOSC2 upon arsenite stress in MCF7 cells (Figure 6B). Under normal conditions EXOCS2 was located in the cytosol as well as the nucleus, changing to pronounced nuclear localization after five minutes of arsenite stress – a situation that persisted throughout the 30 minutes of the experiment. Notably, since our total proteome data had shown that the overall abundance of EXOSC2 (or any other exosomal protein) was not affected by arsenite stress (Table S3), we conclude that EXOSC2 redistributed to the nucleus. Since the change in RNA-binding (Figure 5B) and nuclear localization (Figure 6B) coincided after five minutes of arsenite stress, we hypothesized that EXOSC2 had changed RNA interaction partners in the process. Therefore we performed XRNAX CLIP-seq for EXOSC2 in unstimulated cells, and cells treated with arsenite for five and 30 minutes. Interestingly, we identified particularly nuclear transcripts that increased their association with EXOSC2 upon arsenite stress (Table S5), in addition to a high abundance of tRNA and snoRNA, known to be degraded or processed, respectively, by the exosome in yeast (Gudipati et al., 2012). In addition, on average 65 % (σ=13 %) of unique reads mapped to ribosomal transcripts, in line with another previous study in yeast (Schneider et al., 2012). Since the exosome participates in processing of the 45S pre-rRNA by trimming it into 18S, 5.8S and 28S rRNA (for review see (Henras et al., 2015)), we analysed how read coverage of the 45S pre-rRNA by EXOSC2 changed upon arsenite stress. We found very significant enrichment for specific regions (Figure 6C), most notably in segments 3’ of the mature 5.8S transcript, which is the main region known to be degraded by the exosome during canonical 5.8S rRNA maturation. Interestingly, two peaks demarcating intermediates known as 7S and 6S pre-rRNA (Tafforeau et al., 2013), and a peak 5’ of the mature 18 S transcript, which is known to be cooperatively degraded by the exonuclease XRN2 and the exosome (Sloan et al., 2014), were highly prominent in untreated cells but strongly decreased after 30 minutes of arsenite stress (Figure 6C). Both observations implied that import of EXOSC2 coincided with rRNA processing, eliminating transient intermediates such as the 6S or 7S pre-rRNA. Thus, our data suggested that EXOSC2 was imported into the nucleus upon arsenite stress in order to promote rRNA maturation.

## Discussion

In the past UV-crosslinking has proven an invaluable tool to study protein-RNA interactions *in vitro* and *in vivo* (McHugh et al., 2014), both using sequencing technologies to characterize RNAs that interact with proteins (Lee and Ule, 2018) and, more recently, using mass spectrometry to identify proteins interacting with RNA (Hentze et al., 2018). An invariant requirement for each of these methodologies is the need to enrich for RNA-protein complexes to i) distinguish them from their free constituents, ii) to select for an RNA-species or protein of interest as an entry point, or iii) to increase their abundance and facilitate their detection in the first place. Given this universal need, we have developed XRNAX as a generic method for the biochemical purification of protein-crosslinked RNA. The unique feature of XRNAX is its ability to globally purify protein-crosslinked RNA irrespective of its sequence and biotype (Figure 1C) or the type of UV-crosslinking that is used (Figure 1B). XRNAX is distinct from oligo(dT) interactome capture that primarily targets mRNA (Baltz et al., 2012; Castello et al., 2012), or RBR-ID (He et al., 2016) that relies on crosslinking but does not enrich. Moreover, it is not restricted to nascent RNA as in ‘RNA interactome using click chemistry’ (RICK) (Bao et al., 2018) and presents protein-crosslinked RNA as own entity. We validated the qualitative and quantitative performance of XRNAX by comparing UV-crosslinked to non-crosslinked (Figure 1F), RNA-depleted (Figure S1D) and spike-in controls (Figure S5A), allowing for more stringent cut-offs than previous interactome capture studies and better differential quantification through exclusion of background peptides. These characteristics, combined with the facile interfacing with various transcriptomic and proteomic methodologies position XRNAX for numerous applications in RNA biology.

### The Integrated Human RNA-Binding Proteome Uncovers Novel RNA-Binding Proteins

Our analysis of RNA-bound proteomes confirmed the large majority of the proteins previously shown to interact with poly(A) RNA in the respective cell lines (Figure 3D, Table S2). Beyond this, we identified many hundreds of proteins that we tentatively designated as non-poly(A) interactors because of their prior association with RNA-biology (Figure 3D) while being absent in previous interactome studies, albeit we do not exclude that some of them also bind to mRNA. Within this group of non-poly(A) interactors a striking observation was the enrichment of bromodomain-containing proteins. Chromatin-modifying complexes such as PRC2, CoREST or SMCX have been found to interact with a large number of lincRNAs (Khalil et al., 2009), which in some cases were shown to modulate their function, e.g. the lincRNA HOTAIR recruits PRC2 and leads to repression of genes in the HOXD cluster (Rinn et al., 2007). Future studies should reveal if and how ncRNAs may also regulate the function of bromodomain-containing complexes.

By comparing proteins in the ihRBP we found that specific low-complexity motifs were enriched in poly(A)-binding proteins over non-poly(A)-binding proteins. Growing evidence has linked low-complexity motifs and IDRs in RNA-binding proteins to liquid droplet formation (for review see(Shin and Brangwynne, 2017)). Liquid-liquid phase separation in biological systems is a powerful concept to explain the assembly of biomolecules into functional complexes. Indeed, many of the poly(A)-binding proteins harbouring IDRs, such as TIA, hnRNPA2 or FUS have been implicated in the nucleation of mRNA in macroscopic complexes such as stress granules (Jain et al., 2016; Markmiller et al., 2018), which have been shown to include mRNA and exclude ncRNA (Khong et al., 2017).

### XRNAX Connects Translational Arrest, Stress Granule Formation and Autophagy

When investigating arsenite-induced translational arrest we were surprised to see rapid and specific degradation of the translational machinery (Figure 4C), where cytosolic ribosomal proteins (Figure 4D) and translation initiation factors (Figure S4A) decreased 50% in abundance within 30 minutes. In human cells degradation of translation-associated proteins has been observed upon amino acid starvation (Gretzmeier et al., 2017), however, on a much longer timescale of hours to days instead of minutes. Here we demonstrate that this rapid elimination is the combined effect of reduced protein synthesis (Figure 4A) and autophagy-mediated degradation (Figure 4F), highly reminiscent of a process coined as ribophagy in yeast (Kraft et al., 2008). While a recent report observed protein degradation and induction of autophagy upon arsenite stress in yeast (Guerra-Moreno et al., 2015), these processes were never causally linked or observed in a mammalian system. Our data now showed that the arsenite-induced degradation of translation-associated proteins could be inhibited by spautin-1, indicating that this is driven by autophagy. Interestingly, ribophagy in yeast depends on Ubp3 and Bre5 (Kraft et al., 2008), the orthologues of the human stress granule markers USP10 and G3BP1. This is interesting because we identified USP10 among the proteins with the most prominent decrease in RNA-binding upon arsenite stress (Figure 5E), suggesting that this may be the missing link to explain the previously observed role of USP10 in counteracting arsenite-induced oxidative stress (Takahashi et al., 2013). Notably, USP10 is one of the two known targets of the autophagy inhibitor spautin-1 (Liu et al., 2011), thus, closely tying together USP10 as an RNA-binding protein, a mediator of autophagy, and a constituent of stress granules.

Collectively, our data showed that arsenite simultaneously induced a decrease in protein synthesis and disassembly of polysomes from mRNA, along with selective degradation of the translational machinery. This was concurrent with release of the stress granule protein USP10 from RNA, potentially mediating a multi-faceted and rapid lock on translation.

### XRNAX Opens New Dimensions for Exploring the Protein-RNA Landscape

Although we have shown a range of applications of XRNAX, each one of which has led to novel biological insights, we anticipate that many more can be explored in the future. First, XRNAX is not limited to human cells used here. UV-crosslinking (Darnell, 2010) and TRIZOL extraction have been successfully applied to bacteria, viruses, yeast, plant and animal tissue (Chomczynski and Sacchi, 2006), making them readily accessible for XRNAX. We envision that especially infection biology, involving non-adenylated bacterial and viral RNA, will greatly benefit from our methodology. Second, XRNAX can be interfaced with the wide variety of CLIP-seq-derived methodologies (Lee and Ule, 2018), far beyond the CLIP-method used here (Figure S6A). This should be highly useful for the detailed characterization of RNA-protein interaction sites. Third, XRNAX may be intersected with methods in chemical biology e.g. to selectively isolate RNA (reviewed in (Grozhik and Jaffrey, 2018)) or protein (reviewed in (Chuh et al., 2016)) carrying post-transcriptional or post-translational modifications, respectively. Here the purity of XRNAX extracts allows for experiments, which are impossibly performed from total lysates. This includes enzymatic reactions, where as a proof-of-concept we were able to biotinylate RNA in XRNAX extracts using poly(U) polymerase (data not shown). We envision this to produce innovative and insightful sequencing and MS applications. Fourth, XRNAX offers an efficient and scalable procedure for the extraction of RNA-peptide hybrids. This may spark MS-based strategies for simultaneous sequencing of peptides and RNA in crosslinked hybrids, ultimately describing a global interaction network of the RNA-binding proteome with the protein-bound transcriptome in one experiment (Lenz et al., 2007). Fifth, complementary sampling of the full protein-bound transcriptome (by RNA-seq and XRNAX-RNA-seq, respectively) may be pursued in future applications to allow for much better sequencing depth, and to identify transcripts that change their association with proteins as a consequence of cellular perturbations. Finally, XRNAX should be an excellent starting point to identify proteins that interact with individual RNA species – an application that is usually greatly impaired by genomic DNA and other cellular constituents in crude lysates. Such an approach should reveal valuable mechanistic insights, as recently demonstrated for the lincRNA *Xist* (Chu et al., 2015; McHugh et al., 2015; Minajigi et al., 2015). Facilitated by XRNAX this may be extended to many other lincRNAs whose function remain to be established.

We demonstrated here that XRNAX is a versatile, reproducible and scalable method to give insights into uncharted areas of the transcriptome and the RNA-binding proteome.

## Acknowledgements

We thank Stefan Wilkening (DKFZ Heidelberg) for access to a Covaris ultrasonicator. We thank Sara Fahs for providing potassium acetate and support. We thank Vladimir Benes, Bettina Hase and Nayara Azevedo (Genecore EMBL Heidelberg) for RNA sequencing as well as advice and discussion. We thank Christian Frese, Gianluca Sigismondo and Gertjan Kramer for continuous support and discussion.

## Author contributions

Conceptualization, J.T. and J.K.; Methodology, J.T. and J.K.; Formal Analysis, JT; Formal Analysis of DUF2373, A.P. and A.B.; Formal Analysis of XRNAX-CLIP-seq, T.S.; Investigation, J.T.; Writing – Original Draft, J.T. and J.K.; Writing

– Review & Editing, J.T., M.H. and J.K.; Visualization, J.T.; Supervision, J.K.; Funding Acquisition, J.K.

## Supplementary figure legends

**Figure S1:**
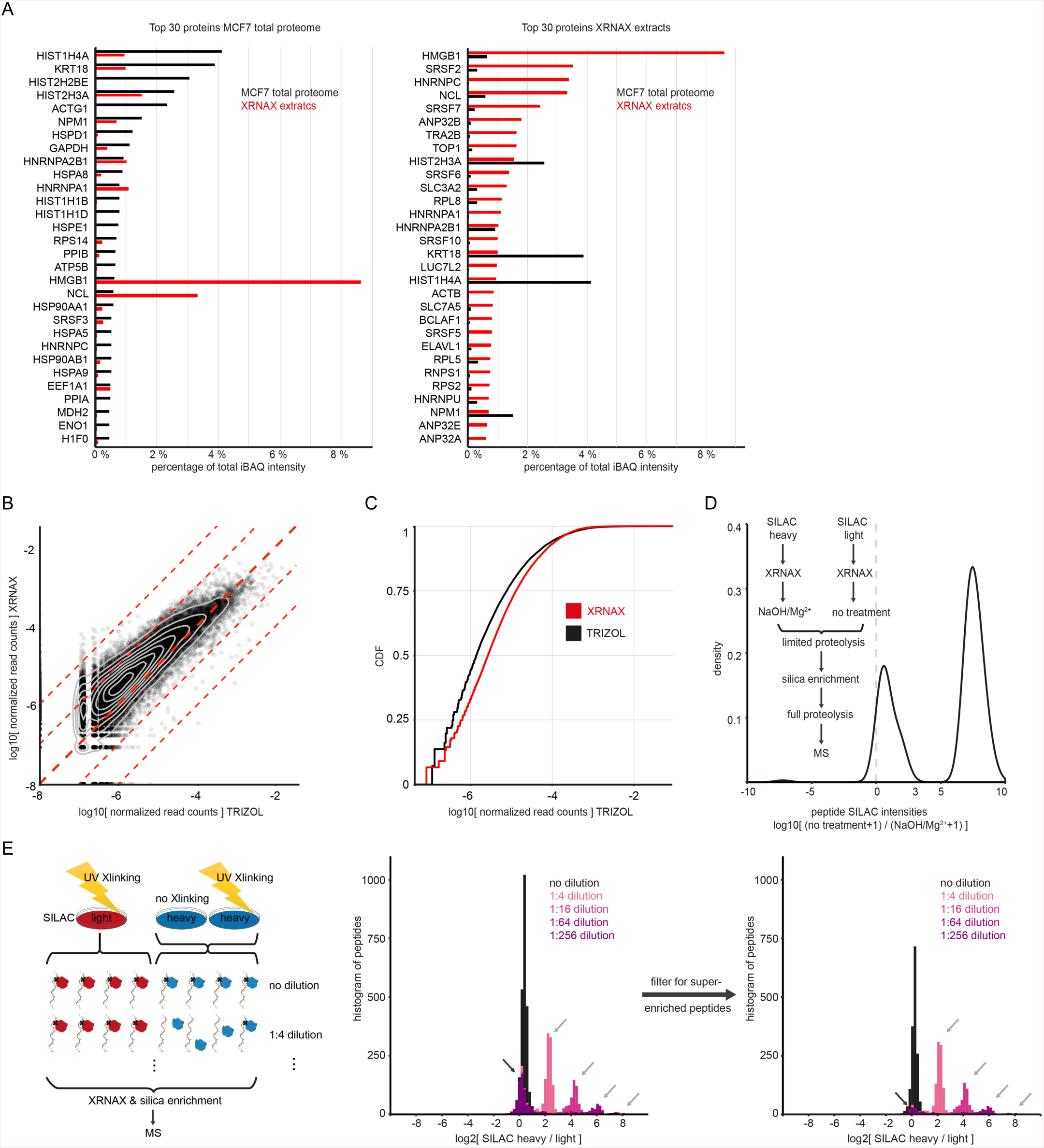
Proteomic and transcriptomic features of XRNAX-extracts. (A) Relative abundance of proteins in the total proteome and XRNAX-extracts of MCF7 cells, estimated from iBAQ values. Bar graphs display contribution of the top-30 proteins to the combined iBAQ intensity of all detected proteins. (B) Scatterplot comparing normalized read counts for all GENCODE-annotated transcripts in RNA obtained by XRNAX and TRIZOL. MCF7 cells were exposed to 4SU for 16 hours before UV-crosslinking at 365 nm and processing via XRNAX, or without crosslinking and conventional TRIZOL extraction. Reads were counted per gene and normalized to the total number of counts. Each point represents one gene and displays the mean of two replicates. Contour lines indicate highest density of the points in the plot. Dashed lines indicate fold-changes of 1, 10 and 100. Sequencing for all replicates was performed in one lane and read count for all libraries was within 10 % deviation from the average read-count. (C) Comparison of normalized read counts for all GENCODE genes between XRNAX and TRIZOL-extracted RNA. Same data as in B shown as cumulative distribution. (D) Density plot showing the enrichment of peptides from XRNAX-extracts with intact RNA over XRNAX-extracts where RNA was degraded. For details see text. (E) Proof-of-concept for the differential quantification of RNA-binding using XRNAX and silica enrichment. Heavy SILAC-labeled MCF7 cells were UV-crosslinked and mixed with non-crosslinked heavy MCF7 cells in 5 defined ratios. These mixtures of heavy cells were combined with the identical amount of UV-crosslinked, light cells and subjected to XRNAX followed by silica enrichment and MS quantification. Histogram displays SILAC ratios without normalization. Peptides that were found super-enriched in previous experiments using a non-UV-crosslinked control (Figure 1F) showed discrete fold-changes corresponding to mixing ratios (grey arrows), whereas peptides that were not super-enriched before showed a 1:1 ratio (black arrow).

**Figure S2:**
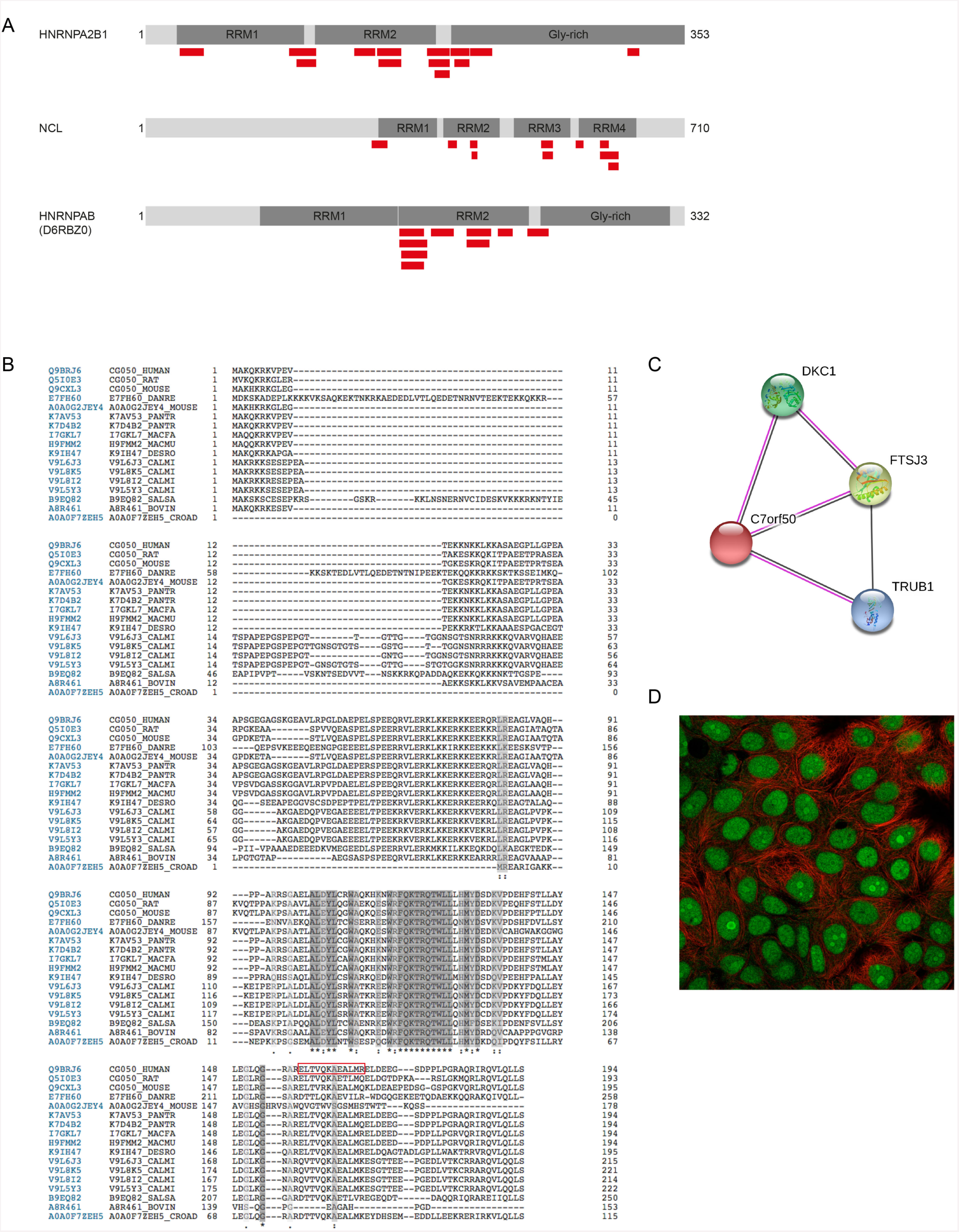
Sequence analysis of C7orf50 and the putative RNA-binding domain DUF2373. (A) Proteins with the largest number of identified cyclic U-crosslinked peptides, indicating their annotated domain structure (dark grey) and the position of the detected cyclic U-containing peptides. (B) Sequence alignment for 17 high-confidence hits from a HMMER search for human C7orf50. The identified cyclic U-crosslinked peptide, ELTVQKAEALMR, is highlighted in red. (C) STRING interaction network for C7orf50. DKC1 and TRUB1 are involved in pseudourylation of rRNA, FTSJ3 in methylation of rRNA. (D) Immunofluorescence image of C7orf50 in MCF7 cells. Red: microtubules; green: C7orf50. Image courtesy: www.proteinatlas.org.

**Figure S3:**
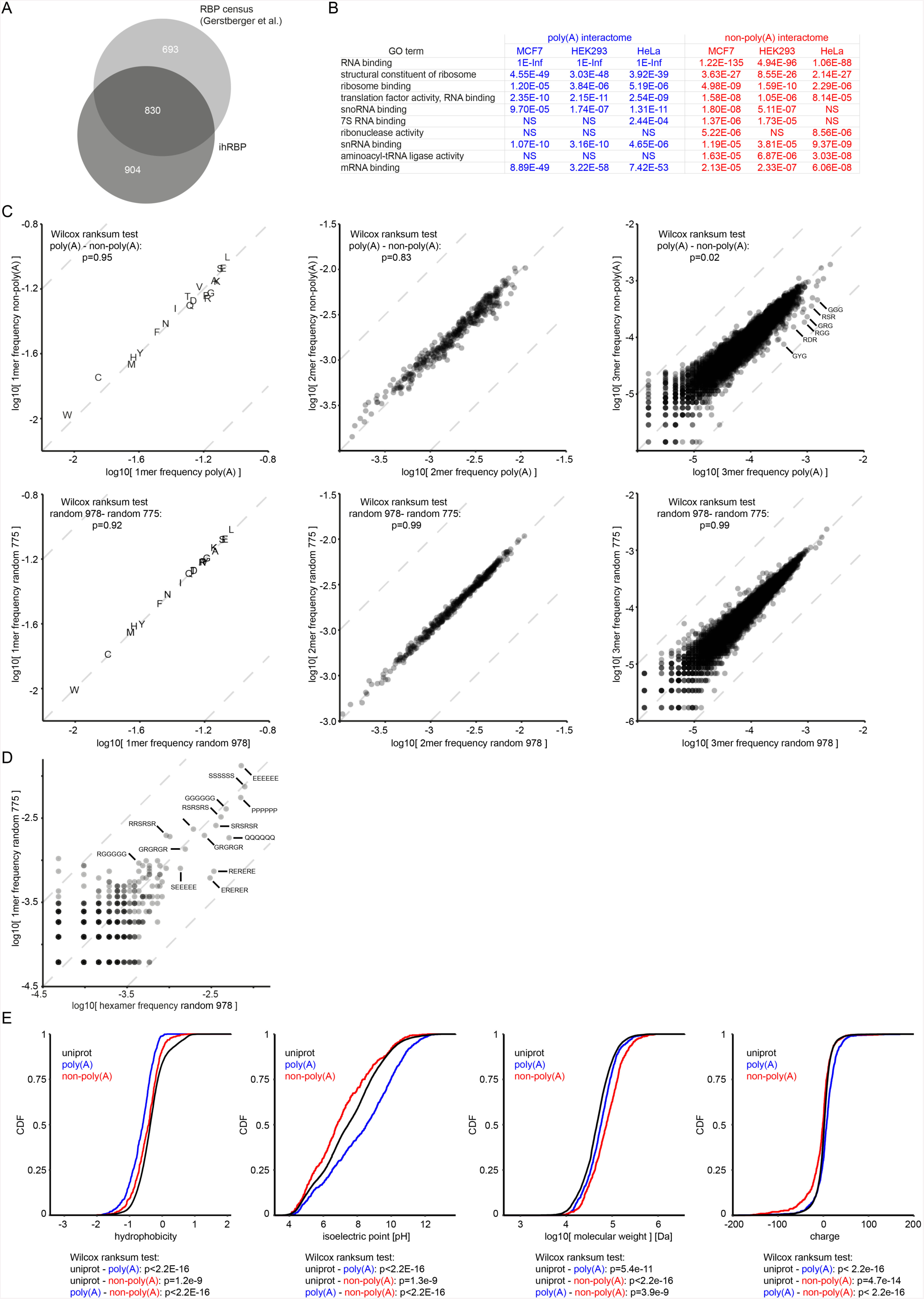
Properties of XRNAX-derived RNA-binding proteomes. (A) Venn diagram comparing proteins of the ihRBP to the census of RNA-binding proteins from Gerstberger et al. (2014). (B) GO enrichment analysis for XRNAX-derived RNA-binding proteomes from three cell lines. Displayed are adjusted p-values for ten RNA-related terms with especially strong enrichment in either group. Note that enrichment analysis was performed against two different background sets, for details see text. Inf: infinity, NS: not significant with a p-value<10E3. (C) Scatter plots comparing amino acid, dipeptide and tripeptide (k-mer) frequenies between poly(A) and non-poly(A) interactomes in the ihRBP (see Figure 3E). Top: All possible permutations of the 20 amino acids for each k-mer were counted in proteins of each group and normalized to the total number of counts. Bottom: Control analysis for two groups of the same size but containing randomly selected proteins from the ihRBP. (D) Control analysis referring to Figure 3F. Scatter plot comparing hexamer frequencies for control groups with randomly selected proteins as described in panel C. (E) Cumulative distributions of physicochemical properties in the poly(A) and non-poly(A) interactomes (see Figure 3E) and the entire UniProt human proteome (uniprot).

**Figure S4:**
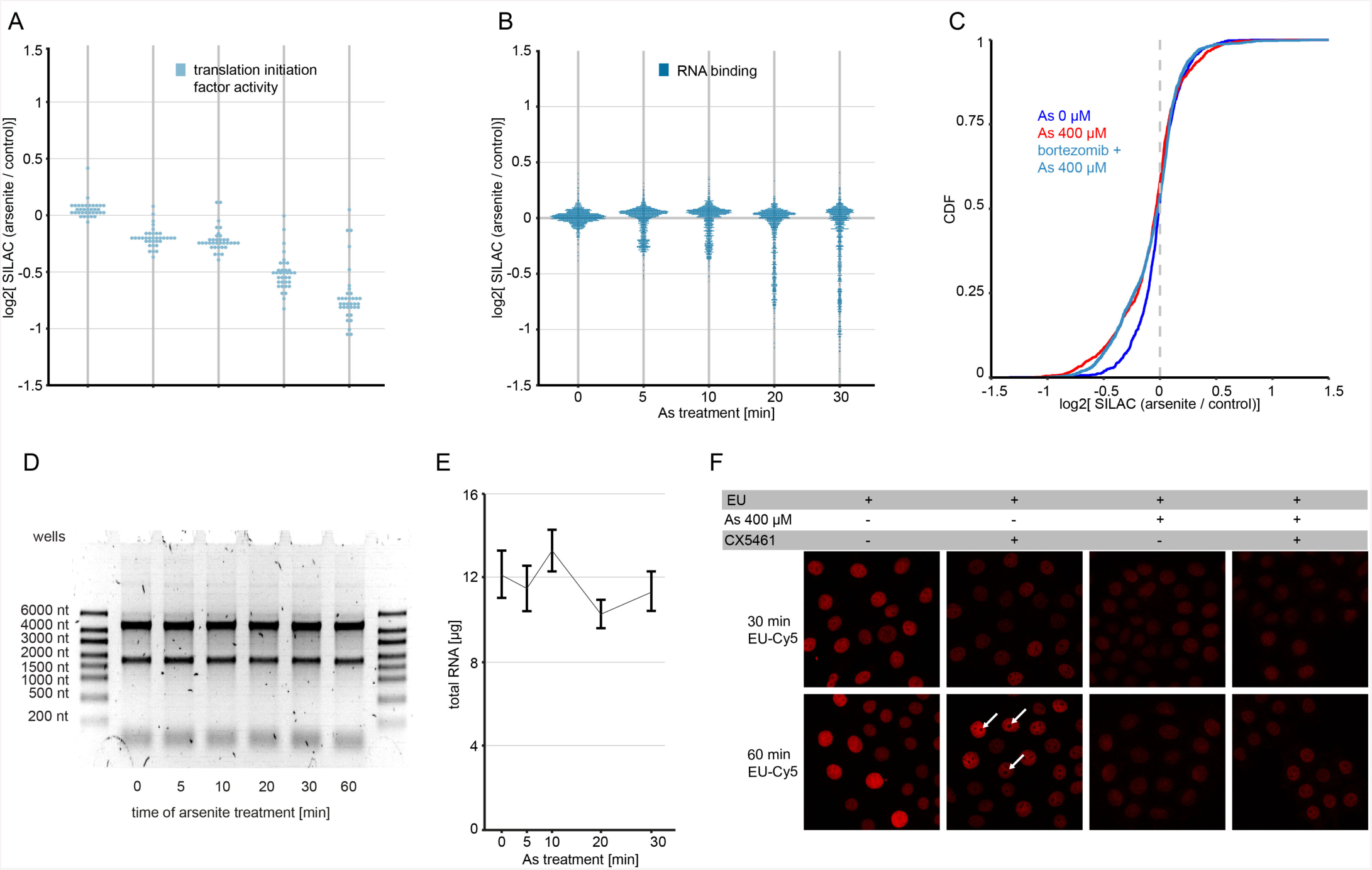
Changes in the total proteome upon arsenite stress. (A) Dotplots displaying changes in total proteome of MCF7 cells upon arsenite stress for proteins with the GO-term ‘Translation initiation factor activity’. Each dot represents one protein. Values displayed are means of duplicate experiments with label swap filtered for a variance of 15 % or smaller. (B) Same as panel A for proteins with the GO-term ‘RNA binding’. (C) Cumulative distribution of changes in total proteome of MCF7 cells upon arsenite treatment and proteasome inhibition by bortezomib. Cells were treated for 1 hour with 500 nM bortezomib (green) or left untreated (red) before heavy SILAC labeled cells were treated with 400 µM arsenite for 30 minutes. (D) Agarose gel electrophoresis of total RNA extracted from MCF7 cells upon 400 µM arsenite stress for indicated time. (D) Timeline showing yield of total RNA extracted from MCF7 cells after arsenite stress. Identical number of cells were treated with 400 µM arsenite for indicated time, total RNA was extracted and quantified using UV-spectrosopy. N=6, error bars indicate standard error of the mean (SEM). (E) Confocal microscopy of MCF7 cells incorporating EU upon arsenite stress. To exclude the possibility that RNA turnover may be increased upon arsenite stress, ethenyl-uridine (EU) labelling was applied to visualize newly synthesized RNA using click-chemistry. After applying the RNA-polymerase I inhibitor CX5461 discrete areas (white arrows), presumably representing nucleoli, were not stained anymore demonstrating specific labelling of nascent transcripts. Under arsenite stress incorporation of EU was lower, indicating reduced transcription. Exchange of cytosolic RNA with nascent RNA was not apparent during the chosen time window.

**Figure S5:**
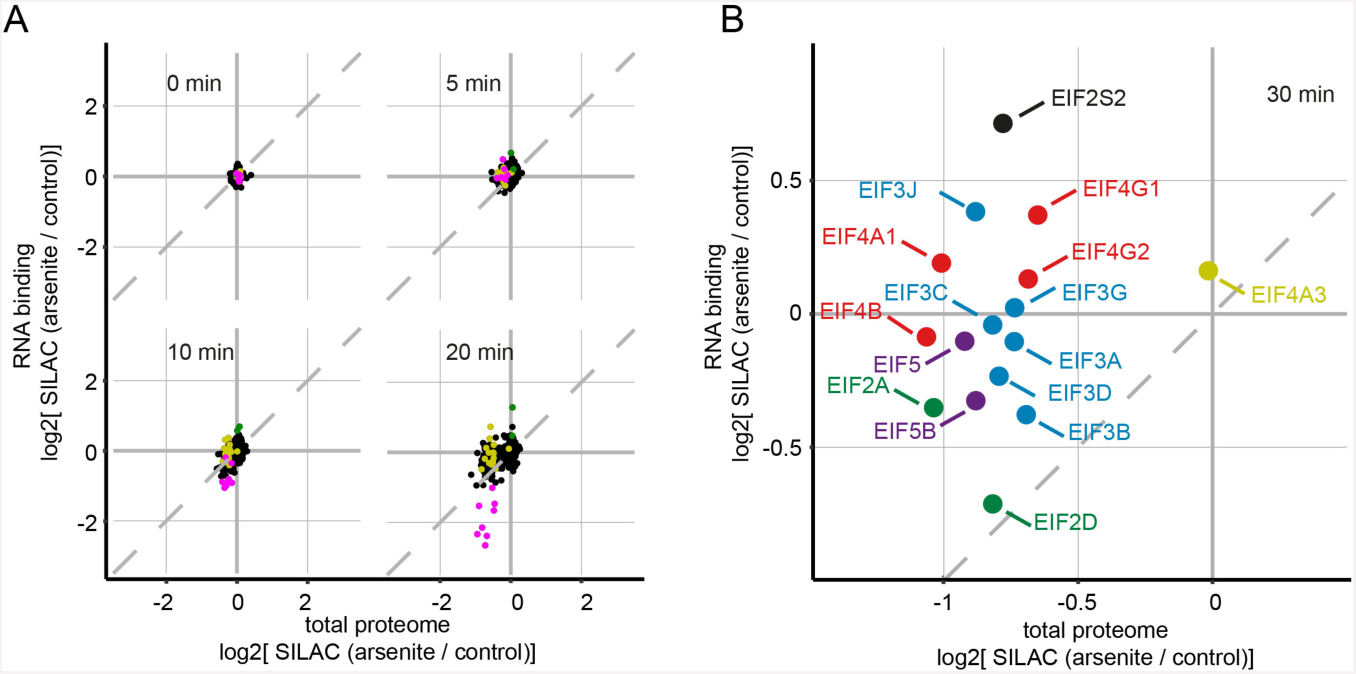
Quantification of changes in RNA-binding proteins. (A) Scatter plots comparing changes in the total proteome to changes in RNA-binding after 0, 5, 10 and 20 minutes of arsenite stress. See also Figure 5. (B) Scatter plot comparing RNA-binding versus total proteome changes after 30 minutes of arsenite treatment. Magnification of data in Figure 5E, only showing eukaryotic translation initiation factors (EIFs). Color coding refers to complexes these proteins are known to be part of. Black: EIF2 complex, blue: EIF3 complex, red: EIF4 complex, magenta: EIF5 complex, yellow: exon junction complex, green: auxiliary EIFs without membership in any core complex.

**Figure S6:**
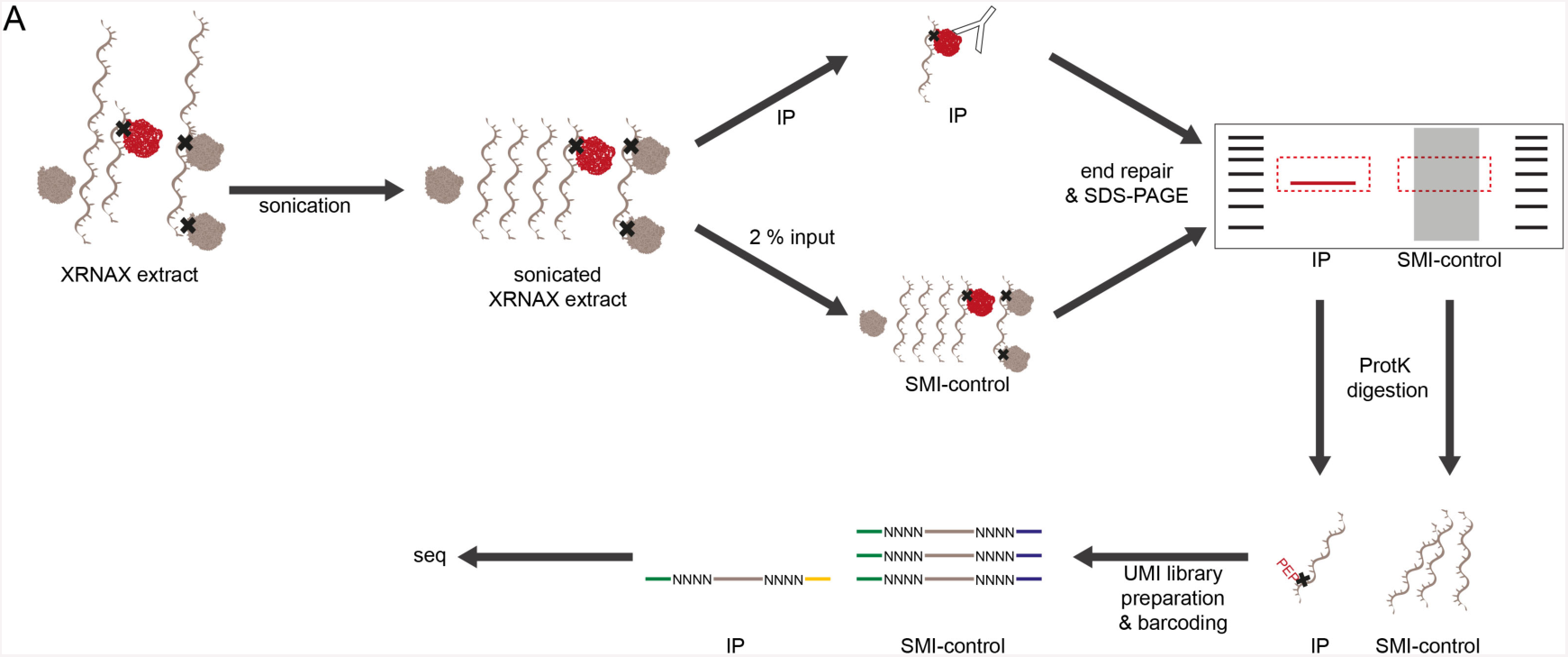
Combination of XRNAX and CLIP-seq. (A) Experimental scheme for an XRNAX CLIP-Seq experiment. After initial fragmentation of RNA through ultrasonication, IP against a protein of interest co-precipitates crosslinked RNA fragments. Both input and IP are resolved on an SDS-PAGE, blotted, and membrane pieces cut out in a region corresponding to the adequate molecular mass. RNA is released by proteinase K digestion and prepared into a sequencing library using conventional small RNA library preparation with unique molecular identifiers (UMIs). For further details refer to method section.

## Supplemental information

**Table S1:** Cyclic Uridine Monophosphate-Crosslinked Peptides Identified by MS

**Table S2**: RNA-Binding Proteomes of MCF7, HEK293 and HeLa Cells

**Table S3**: Changes Upon Arsenite Stress Quantification of the MCF7 Total Proteome

**Table S4**: Changes Upon Arsenite Stress Quantification of the MCF7 RNA-Interactome

**Table S5**: XRNAX-CLIP-seq Non-Ribosomal Transcripts Interacting with EXOSC2

### KEY RESOURCES TABLE

| REAGENT or RESOURCE | SOURCE | IDENTIFIER |
| --- | --- | --- |
| Antibodies |  |  |
| Lamin B1 rabbit polyclonal antibody | proteintech | 12987-1-AP,<br>RRID:AB_2136290 |
| EXOSC2 rabbit polyclonal antibody | proteintech | 14805-1-AP,<br>RRID:AB_2101837 |
| EXOSC2 mouse monoclonal antibody | proteintech | 66099-1-Ig |
| Goat anti-mouse Cy5 | abcam | ab6563,<br>RRID:AB_955068 |
| Chemicals, Peptides, and Recombinant Proteins |  |  |
| Dialysed FBS | Gibco | 26400-044 |
| Pen-Strep | Gibco | 15140-122 |
| DMEM for SILAC | Silantes | 280001300 |
| SILAC heavy L-lysine ($^{13}\text{C}_6$, $^{15}\text{N}_2$-L-Lysine HCl) | Silantes | 211604102 |
| SILAC heavy L-arginine ($^{13}\text{C}_6$, $^{15}\text{N}_4$-L-Arginine HCl) | Silantes | 201604102 |
| GlutaMAX | Gibco | 35050061 |
| TRI reagent | Sigma | T9424 |
| GlycoBlue | Ambion | AM9515 |
| 4-thiouridine (4SU) | biomol | Cay-16373 |
| SYBRSafe | invitrogen | S33102 |
| EDTA-free protease inhibitor | Sigma (Roche) | 11873580001 |
| Benzonase | Novagen | 70664 |
| SP3 beads | GE | 44152105050250 |
| Trypsin/LysC | Promega | V5073 |
| NEB DNase buffer 10 x | NEB | B0303S |
| NEB DNase | NEB | M0303L |
| RNASin Plus RNase inhibitor | Promega | N2615 |
| RNase A | Thermo | EN0531 |
| RNase I | Ambion | AM2295 |
| RNase T1 | Thermo | EN0541 |
| Sodium arsenite (50 mM solution in water) | Santa Cruz | sc-301816 |
| Spautin-1 | Sigma | SML0440 |
| Rapamycin (2.5 mg/ml in DMSO) | Sigma | R8781 |
| Ethynyl-uridine (EU) | Jena Biosciences | CLK-N002 |
| RNA polymerase I inhibitor CX5461 | Millipore | 509265 |
| sulfo-Cy5-azide | Jena Biosciences | CLK-AZ118 |
| HOECHST33342 (20 mM in water) | Thermo | 62249 |
| ProlongGold antifade mountant | invitrogen | P36934 |
| Proteinase K | Thermo | EO0491 |
| FastAP Thermosensitive Alkaline Phosphatase | Thermo | EF0651 |
| T4 Polynucleotide Kinase | Thermo | EK0032 |
| Protein G Magnetic Beads | Pierce | 88847 |
| Critical Commercial Assays |  |  |
| Quiagen RNeasy Midi Kit | Quiagen | 75144 |
| NEXTflex Small RNA Sequencing Kit | Bioo Scientific | NOVA-5132 |
| Deposited Data |  |  |
| currently uploading |  |  |
| Experimental Models: Cell Lines |  |  |
| Human: MCF7 cells | ATCC | RRID:CVCL_0031 |
| Human: HEK293 cells | Laboratory of Rolf Sprengel (MPI Heidelberg) | RRID:CVCL_0045 |
| Human: HeLa cells | ATCC | RRID:CVCL_0030 |
| Software and Algorithms |  |  |
| GOrilla | (Eden et al., 2009) | http://cbl-gorilla.cs.technion.ac.il/ |
| MEME | (Bailey et al., 2009) | http://meme-suite.org/index.html, RRID:SCR_001783 |
| BBMap (37.68) |  | http://sourceforge.net/projects/bbmap |
| cutadapt (version 1.10) | (Martin, 2011) | http://cutadapt.readthedocs.io/en/stable/index.html, RRID:SCR_011841 |
| Je (version 1.2) | (Girardot et al., 2016) | https://git.embl.de/grp-gbcs/Je |
| STAR (version 2.5.0a) | (Dobin et al., 2013) | https://github.com/alexdobin/STAR, RRID:SCR_015899 |
| Bowtie2 | (Langmead and Salzberg, 2012) | http://bowtie-bio.sourceforge.net/bowtie2/index.shtml |
| HTSeq | (Anders et al., 2015) | https://htseq.readthedocs.io/en/release_0.9.1/, RRID:SCR_005514 |
| UCSF Chimera (1.12) | (Pettersen et al., 2004) | https://www.cgl.ucsf.edu/chimera/, RRID:SCR_004097 |
| MaxQuant (1.5.1.2) | (Cox and Mann, 2008) | http://www.biochem.mpg.de/5111795/maxquant, RRID:SCR_014485 |
| MSFragger | (Kong et al., 2017) | http://www.nesvilab.org/software.html |
| R | R Core Team (2016). R: A language and environment for statistical computing. R Foundation for Statistical Computing, Vienna, Austria. | https://www.r-project.org/, RRID:SCR_001905 |
| RStudio (0.99.903, RRID:SCR_000432) | RStudio: Integrated Development for R. RStudio, Inc., Boston, MA | http://www.rstudio.com/,<br>RRID:SCR_000432 |
| Peptides (R) |  | https://github.com/dosorio/Peptides/ |
| ggplot2 (R) |  | http://ggplot2.tidyverse.org/,<br>RRID:SCR_014601 |
| Biostrings (R) |  | http://bioconductor.org/packages/Biostrings/ |
| csaw (R) | (Lun and Smyth, 2015) | http://bioconductor.org/packages/csaw/ |
| DESeq2 (R) | (Love et al., 2014) | http://bioconductor.org/packages/DESeq2/,<br>RRID:SCR_015687 |
| Other |  |  |
| Advanced online documentation for XRNAX protocols |  | https://www.xrnax.com/ |

## Experimental Model and Subject Details

### Mammalian Cell Culture and Stable Cell Lines

All cell lines were maintained in Dulbecco’s Modified Eagle’s Medium (DMEM) for SILAC supplemented with 10% dialysed FBS and Pen-Strep (100 U/ml penicillin, 100 mg/ml streptomycin) at 37 °C, 5 % CO_2_. DMEM for SILAC was supplemented with 1 mM L-lysine and 0.5 mM L-arginine of the individual SILAC labels as well as 1.7 mM light L-proline and 1 x GlutaMAX. The heavy SILAC label was introduced during six passages in heavy DMEM for SILAC.

## Method Details

### Advanced Online Documentation

In order to make XRNAX and its applications accessible to a wide audience and to promote the development of second party applications we created a website accessible under www.xrnax.com, where detailed protocols are presented with schemes and illustrations. (Currently password protected).

### Guanidinium Thiocyanate–Phenol–Chloroform (TRIZOL) Extraction

Up to 10 million cells were lysed in 1 ml TRI reagent by pipetting up and down. For phase-separation, 200 µl chloroform was added and samples mixed by turning tubes upside down several times. After 5 min incubation at room temperature, samples were spun down with 12000 g for 10 min at 4 °C. Approx. 400 µl of the aqueous phase was transferred to a fresh tube, NaCl was added to a final concentration of 300 mM along with 1 µl GlycoBlue. Samples were combined with 500 µl isopropanol, mixed by inversion and RNA precipitated by centrifugation with 18000 g for 15 min at-10 °C.

The supernatant was removed and the RNA pellet washed with 1 ml of 70 % ethanol before resuspension in desired volume of nuclease-free water.

### UV-Crosslinking of Cells

Cells were grown in 245 mm x 245 mm dishes to desired confluence. For the incorporation of 4-thiouridine (4SU) into RNA, cells were incubated with 100 µM 4SU for 16 hours prior to UV-crosslinking. Media was decanted and cells washed with 50 ml ice-cold PBS. In order to remove as much liquid as possible dishes were propped up straight and residual PBS drained onto a paper towel through gravity. UV-crosslinking occurred on ice with 200 mJ/cm^2^ at 254 nm wavelength with a BIO-LINK UV-crosslinker (Vilber). Cells that had incorporated 4SU were UV-crosslinked at 365 nm wavelength. Subsequently, cells were harvested into ice-cold PBS, pelleted and either directly subjected to XRNAX or stored at −80 °C for up to 14 days.

### RNA Analysis Using Agarose Gel-Electrophoresis

To verify the integrity of RNA extracted by TRIZOL or XRNAX (see below), agarose-gel electrophoresis was performed using 1 % agarose in TBE and SYBRSafe staining. Specifically, for Figure 1B 0.05 % of the total yield extracted from 10 million MCF7 cells using the indicated method was subjected to the indicated treatment. Samples were denatured in RNA gel loading dye containing formamide for 2 minutes at 85 °C and run for 40 minutes with 3 W.

### Proteomic Sample Preparation

For MS sample preparation, a modification of the SP3 protocol described by Hughes et al. (Hughes et al., 2014) was used. For total proteome analysis approx. 1 million cells were lysed and reduced in 1 ml lysis buffer (Tris-Cl 50 mM, DTT 10 mM, SDS 0.05%) at 95 °C, 700 rpm shaking for 30 minutes. For samples other than cells, e.g XRNAX extracts, samples were brought to a total volume of 100 µl with MilliQ water and combined with 900 µl lysis buffer before reduction at 95 °C, 700 rpm shaking for 30 minutes. Magnesium chloride (final concentration of 5 mM), CAA (20 mM), and EDTA-free protease inhibitor were added and mixed before addition of 1 µl of benzonase. Subsequently, digestion of nucleic acids and alkylation occurred for 2 hours at 37 °C, 700 rpm shaking. 400 µl SP3 beads were preconditioned by washing with MilliQ water 3 times, before reconstitution in 1 ml MilliQ water. EDTA was added to 10 mM final concentration along with 1 % SDS and 20 µl SP3 beads. Samples were vortexed vigorously and subsequently combined with 1 ml acetonitrile. Samples were mixed again and incubated for 15 minutes at room temperature for protein binding to occur. The beads were collected on a magnetic stand for 2 min and supernatants decanted. While in the magnetic stand beads were then washed 3 times with 2 ml ethanol 70 %, which was added for 1 minute and subsequently decanted. Residual ethanol was removed and beads were taken up in the desired digestion volume of TEAB 20 mM and adequate amounts of trypsin/LysC added to the solution (for total proteomes from 1 million cells 1 µg trypsin/LysC in 100 µl TEAB). Samples were digested at 37 °C, 700 rpm shaking overnight. For single run analysis formic acid was added to a final concentration of 1 % and samples spun down for 5 minutes with 20000 g. Supernatants were transferred to fresh tubes without disturbing the pellet and analysed by HPLC-MS.

High pH reversed-phase fractionation occurred under standard settings described below. Of the 40 collected fractions the initial 8 fractions up to approx. 18 % B were discarded, the following 32 fractions were combined to 8 using the scheme 1+9+17+25/…/8+16+24+32. The combined fractions were dried by SpeedVac and taken up in 1 % formic acid before analysis by HPLC-MS.

### High pH Reversed-Phase Fractionation

Fractionation at high pH occurred on an Agilent Infinity 1260 LC system (Agilent) using a Phenomenex Gemini 3 µM C18, 100 x 1 mm column (Phenomonex). Buffer A was NH_4_COOH 20 mM, buffer B was 100 % acetonitrile. The following gradient was used for all applications described in this manuscript: 0-2 minutes 0 % B, 2-60 minutes linear gradient to 65 % B, 61-62 minutes linear gradient to 85 % B, 62-67 minutes 85 % B, 67-85 minutes 0 % B. Eluates were collected in 40 fractions and combined as described in the individual paragraphs.

### HPLC-MS

Separation by HPLC prior to MS occurred on an Easy-nLC1200 system (Thermo Scientific) using an Acclaim PepMap RSCL 2 µM C18, 75 µm x 50 cm column (Thermo Scientific) heated to 45 °C with a MonoSLEEVE column oven (Analytical Sales and Services). Buffer A was 0.1 % formic acid, buffer B was 0.1 % formic acid in 80 % acetonitrile. The following gradient was used for all applications described in this manuscript: 0 minutes 3% B, 0-4 minutes linear gradient to 8 % B, 4-6 minutes linear gradient to 10 % B, 6-74 minutes linear gradient to 32 % B, 74-86 minutes linear gradient to 50 % B, 86-87 minutes linear gradient to 100 % B, 87-94 minutes 100 % B, 94-95 linear gradient to 3 % B, 95-105 minutes 3 % B.

Single-run total proteome analysis was performed on a Fusion Orbitrap mass spectrometer (Thermo Scientific). MS1 detection occurred in orbitrap mode at 60000 resolution, AGC target 1E6, maximal injection time 50 ms and a scan range of 375-1500 DA. MS2 detection occurred with an HCD collision energy of 33 in ion trap top20 mode with an isolation window of 1.6 Da, AGC target 1E4 and maximal injection time of 50 ms.

Detection of XRNAX-derived nucleotide-crosslinked peptides, XRNAX-derived RNA-binding proteomes and XRNAX-derived differential analysis of RNA-binding upon arsenite stress, as well as all analysis of fractionated total proteome samples was performed on a QExactive HF mass spectrometer (Thermo Scientific). MS1 detection occurred at 120000 resolution, AGC target 3E6, maximal injection time 32 ms and a scan range of 350-1500 DA. MS2 occurred with stepped NCE 26 and detection in top20 mode with an isolation window of 2 Da, AGC target 1E5 and maximal injection time of 50 ms.

### Quantification of Nascent Protein Upon Arsenite Stress Using AHA-Labeling

MCF7 cells with heavy and light SILAC labels were expanded for three days on 15 cm dishes until 80 % confluency. Cells of one SILAC label were treated with 400 µM sodium arsenite for 5, 10, 20, 30 or 60 minutes, while cells of the complementary label were left untreated. AHA-labeling and protein purification using click-chemistry for MS quantification was performed as described before (Eichelbaum et al., 2012). In brief, cells were deprived of methionine using methionine-free media for 30 minutes. Labeling started with the addition of AHA-containing media, which occurred simultaneously with the addition of arsenite. After labeling, cells were immediately transferred onto ice, washed with ice-cold PBS and harvested by scraping. Protein enrichment was performed using the Click-iT Protein Enrichment Kit (Thermo Scientific) using the manufacturer’s instructions. Protein captured on agarose beads was subjected to tryptic digestion using 500 ng trypsin/LysC in 200 µl TEAB 50 mM and peptides were cleaned up using an Oasis PRiME HKB µElution Plate (Waters). HPLC-MS detection occurred on an Orbitrap Fusion MS using the parameters described above.

### Protein-Crosslinked RNA Extraction (XRNAX)

Up to 100 million cells (typically one confluent 245 mm x 245 mm dish of UV-crosslinked cells combined with one confluent 245 mm x 245 mm dish of non-crosslinked cells) were lysed in 8 ml TRI reagent by pipetting up and down. Cell clumps were disintegrated by flushing the lysate repeatedly against the wall of the tube. Lysis was further facilitated by incubation on a rotating wheel for 5 minutes at room temperature. Lysates were combined with 1.6 ml chloroform, mixed by inversion and incubated for 5 minutes at room temperature. Tubes were spun down with 7000 g for 10 minutes at 4 °C.

The aqueous phase was removed and the interphase transferred to a 2 ml tube. The interphase was gently washed twice with 1 ml low SDS buffer (Tris-Cl 50 mM, EDTA 1 mM, SDS 0.1 %), flushing protein off the walls of the tube while retaining the integrity of the interphase flakes. Flakes were spun down with 5000 g for 2 min at room temperature and the supernatant discarded. After the washing, flakes were disintegrated by pipetting into 1 ml of low SDS buffer. The disintergrated interphase was spun down with 5000 g for 2 min at room temperature and the supernatant saved as interphase eluate 1. Disintegration of the interphase was repeated with another 1 ml of low SDS buffer, then twice with 1 ml of high SDS buffer (Tris-Cl 50 mM, EDTA 1 mM, SDS 0.5 %) each time yielding approx. 1 ml of interphase eluates. NaCl was added to a final concentration of 300 mM to each of the 4 interphase eluates, along with 1 µl GlycoBlue and 1 ml isopropanol before mixing by inversion. Samples were spun down for 15 min with 18000 g at −10 °C. The supernatants were discarded and pellets from all four elutes were combined in 2 ml of 70 % ethanol. The combined sample was again centrifuged for 1 min with 18000 g at room temperature, supernatant discarded and all residual ethanol removed. The pellet was taken up in 1.8 ml of nuclease-free water and detached from the wall of the tube with a pipette tip. The pellet was allowed to swell for 1 hour on ice with occasional mixing by inversion and eventually dissolved by pipetting.

200 µl NEB DNase I buffer 10 x was added along with 2 µl RNasin Plus, 100 µl NEB DNase and incubated for 60 minutes at 37 °C and 700 rpm shaking. Subsequently, the sample was isopropanol precipitated as described above without further addition of GlycoBlue. Pellets were taken up in 1000 µl nuclease-free water and dissolved by pipetting. RNA concentration was estimated by UV-spectroscopy on a NanoDrop One UV photospectrometer (Thermo Scientific), neglegting adsorbtion by protein.

Purification of protein-free RNA from XRNAX extracts after proteinase K digestion showed that this estimation was within 15 % of the actual RNA content. All amounts of XRNAX extracts mentioned in the following are given in µg of RNA referring to this estimation and do not take protein content into account.

For a detailed, photo-documented version of the XRNAX protocol visit www.XRNAX.com/theprotocol.

### Isolation of Nucleotide-Crosslinked Peptides from XRNAX Extracts

For the isolation of nucleotide-crosslinked peptides, 1000 µg of XRNAX extract were produced from MCF7 cells using the extraction method described above (from 2 confluent 245 mm x 245 mm dishes). Two aliquots of 500 µg of XRNAX extract were brought to 950 µl final volume containing 50 mM Tris-Cl, 0.1 % SDS and 10 mM DTT. 10 µg trypsin/LysC was added to each aliquot to a final volume of 1 ml and digestion occurred for 1 hour at 37 °C, 700 rpm shaking. CAA was added to a final concentration of 20 mM and digestion continued for another hour. Purification of peptide-crosslinked RNA from the digests occurred by silica column purification using the Qiagen RNeasy Midi Kit with modified protocol (refer to kit manual for buffer descriptors). 1 ml digest was combined with 3.5 ml buffer RLT in a 15 ml falcon tube, mixed by inversion and heated to 60 °C for 15 min. The sample was allowed to reach room temperature. 2.5 ml of 100 % ethanol was added, the sample mixed by inversion and applied to an RNeasy Midi column by centrifugation with 3000 g for 5 minutes. Washing occurred twice with 2.5 ml buffer RPE, buffer RW1 was not used. Elution occurred twice with 250 µl nuclease-free water. All eluates combined to approx. 900 µl, which were transferred to a fresh tube. NaCl was added to a final concentration of 300 mM along with 1 µl glycoblue, 1 ml isopropanol, the sample mixed by inversion and incubated for 1 hour at −20 °C. Precipitation occurred by centrifugation with 18000 g at −10 °C for 60 minutes. The supernatant was discarded and the pellet washed with 70 % ethanol. All residual ethanol was removed and the pellet taken up in 60 µl tris-Cl 10 mM. The sample was heated to 85°C for 5 minutes and cooled on ice before addition of 1.5 µl of RNase A, RNase I and RNase T1. RNA digestion occurred for 12 hours at 37 °C, 700 rpm shaking before the sample was heated to 85 °C again for 5 minutes and cooled on ice. Another 1.5 µl of RNase A, RNase I and RNase T1 was added and the sample digested for another 12 hours. High pH reversed-phase fractionation occurred under standard settings described above. The initial peak with high adsorption up to approx. 18 % B containing RNA contaminations was discarded, the following fractions combined, completely dried by SpeedVac, taken up in 1 % formic acid and analyzed by HPLC-MS.

For a detailed version of this protocol and technical advice visit www.xrnax.com/applications.

### SILAC-Controlled Discovery of RNA-Binding Proteins From XRNAX Extracts

To maximize coverage of the RNA-bound proteome, we produced XRNAX extracts from half-confluent and confluent cells (∼40 million cells per condition), each of which were subjected to silica purification after 15 or 30 min of partial tryptic digestion. Cells of one SILAC label were crosslinked with UV-light of 254 nm wavelength as described above, while control cells of the complementary label stayed non-crosslinked. Crosslinked and non-crosslinked cells were combined and extracted by XRNAX as described above.

Per replicate, 930 µg of XRNAX extract (in 930 µl) was further processed. Therefore tris-Cl was added to a final concentration of 50 mM, DTT to 10 mM and SDS to 0.1 % before 20 minutes of incubation at 60 °C, 700 rpm shaking. CAA was added to a concentration of 20 mM and samples incubated for 20 minutes at room temperature. For predigestion, 100 ng trypsin/lysC was added and samples were pre-digested at 37 °C, 700 rpm shaking for 15 or 30 minutes, respectively.

Purification of protein-crosslinked RNA from the digests occurred by silica column purification using the Qiagen RNeasy Midi Kit with modified protocol (refer to kit manual for buffer descriptors). Predigestion was stopped by combining the sample (approx. 1 ml) with 3.5 ml buffer RLT. The sample was mixed by inversion and heated to 60 °C for 15 min. 2.5 ml of 100 % ethanol was added, the sample mixed by inversion and applied to an RNeasy Midi column by centrifugation with 3000 g for 5 minutes. The flow-through was saved for additional rounds of purification. Washing occurred twice with 2.5 ml buffer RPE. Elution occurred twice with 250 µl nuclease-free water. The purification was repeated 3 times, each time using the saved flow through and reusing the same RNeasy Midi column for the individual sample. To the combined eluates NaCl was added to a final concentration of 300 mM along with 1 µl glycoblue and 1 ml isopropanol. The sample was mixed by inversion and incubated for 1 hour at −20 °C. Precipitation occurred by centrifugation with 18000 g at −11 °C for 30 minutes. The supernatant was discarded and the two pellet washed with 70 % ethanol and taken up in 65 µl tris-Cl 50 mM. The sample was heated to 85°C for 5 minutes and cooled on ice before 2.5 µl of RNase A, RNase I and RNase T1 was added. RNA digestion occurred over night at 37 °C, 700 rpm shaking. 500 ng trypsin/LysC were added for 16 hours digestion at 37 °C, 700 rpm shaking. High pH reversed-phase fractionation occurred under standard settings described above. The initial fractions up to approx. 18 % B containing RNA contaminations and few peptides were discarded, the following eluate was collected in six consecutive fractions, which were subsequently dried by SpeedVac and taken up in 1 % formic acid before analysis by HPLC-MS.

For a detailed version of this protocol and technical advice visit www.xrnax.com/applications.

### Differential Quantification of RNA-Binding Upon Arsenite Stress

For the differential quantification of RNA-binding upon arsenite stress, MCF7 cells were expanded for three days to approx. 70 % confluency. 30 million cells of one SILAC-label were exposed to 100 µM sodium arsenite for 0, 5, 10, 20, or 30 minutes, while control cells of the complementary label remained untreated. Duplicate experiments were performed for each time point, which included SILAC-label swap. Both treated and control cells were UV-crosslinked, combined and subjected to XRNAX and silica-enrichment as described above with the only difference that the predigestion time was kept constant at 30 min for all samples. Samples were high pH reversed-phase fractionated into 8 fractions as described above.

Total proteomes where analyzed from cells treated with arsenite at the identical time points, in duplicates and with SILAC label-swap. After treatment the media was discarded and cells were immediately put on ice and washed with ice-cold PBS. Cells were harvested by scraping and subjected to the standard proteomic workflow described above before fractionation into 8 fractions at high pH.

For a detailed version of this protocol and technical advice visit www.xrnax.com/applications.

### Total Proteome Analysis of Arsenite-Induced Protein Degradation

For total proteome analysis of MCF7 under controlled cell culture conditions 0.5 x 10^6^ cells were seeded in 10 cm dishes and cultured for 3 days. Inhibition of autophagy through 10 µM spautin-1 was induced 24 hours prior to arsenite stress. Sodium arsenite was applied at 400 µM concentration. Cells were harvested and subjected to the standard proteomic workflow described above.

We note here that the degree to which autophagic degradation was induced heavily depended on the growth state that the cells were in: During the initial lag-phase of cell culture one day after seeding, arsenite-induced protein degradation was minor compared to the effect observed in the following days of culture (data not shown). For cells seeded at a density to reach confluence after 5 days of culture the most pronounced effect was observed after 3 days.

### MS Database Search

All MS raw files were searched using MaxQuant, except for data of nucleotide-crosslinked peptides. The database searched was the reviewed UniProt human proteome (search term: ‘reviewed:yes AND proteome:up000005640’, 20216 entries, retrieved 11 September 2017) and the default Andromeda list of contaminants. All settings were used at their default value, except for specifying SILAC configurations and indicating the appropriate number of fractions per sample. For the differential quantification of RNA-binding during arsenite stress the match-between-runs option was activated, for all other searches this was explicitly not the case.

MS data of nucleotide-crosslinked peptides was searched with MSFragger using the same UniProt database as mentioned above. Precursor mass tolerance was set to 1000 Da and the export format set to tsv, otherwise all settings were used at their default value.

### Processing and Analysis of MS Data

Gene ontology (GO) enrichment analysis was performed with GOrilla, using either the background-controlled or rank-based mode as detailed in the text. Nucleotide-crosslinked peptides from ribosomal proteins and ribosomal proteins affected in their RNA-binding upon arsenite stress were located in the crystal structure of the human ribosome (Khatter et al., 2015) (PDB 4UG0) using UCSF Chimera.

For the analysis of RNA-binding proteins from XRNAX extracts, the MaxQuant peptides.txt table was filtered to remove entries in ‘potential contaminants’ and ‘reverse’. Furthermore only peptides that matched the category ‘Unique Groups’ were used. To derive RNA-binding proteins for the individual cell lines, peptides from the four replicates were combined and filtered with the condition (SILAC intensity crosslinked +1) / (SILAC intensity non-crosslinked +1)>1000. Pseudo-counts were added to include peptides where the non-crosslinked channel had zero intensity. Proteins (’Leading razor protein’) identified with two or more unique peptides were included in the RNA-binding proteome of the individual cell line.

K-mer frequencies in proteins of the ihRBP were determined using the R package biostrings. Global protein sequence features were computed using the R package ‘peptides’ using the scales ‘Kyte-Doolittle’ for hydrophobicity, ‘EMBOSS’ for isoelectric point and ‘EMBOSS’ for charge.

For differential quantification of RNA-binding upon arsenite stress, the MaxQuant peptides.txt table was filtered to remove entries in ‘potential contaminants’ and ‘reverse’. Furthermore only peptides that matched the category ‘Unique Groups’ and which occurred in the list of ihRBP super-enriched peptides were used. For the quantification of individual proteins (’Leading razor protein’) that were quantified with more than two peptides over all time points the median of normalized SILAC Ratios was computed. Control total proteome data for the experiment was analyzed identically, except for filtering for ihRBP super-enriched peptides, which was omitted. The combined data presented in this manuscript was the mean of biological replicates for each time point filtered for a variance of 15 % or smaller.

For the analysis of the arsenite-induced degradation normalized ratios from the MaxQuant proteinGroups.txt table were used. Dose-dependence was analysed using the R package ‘drc’.

### Ethynyl-Uridine Incorporation and Confocal Microscopy

For the visualization of nascent transcripts using ethynyl-uridine (EU), MCF7 cells were grown on glass cover slips for 3 days. EU was applied at 1 mM, sodium arsenite at 400 µM and the RNA polymerase I inhibitor CX5461 at 10 µM concentration. Treatment occurred for 30 or 60 minutes, the media was discarded and cells washed once with PBS. Fixation occurred with 3 % paraformaldehyde in PBS at room temperature for 10 minutes subsequent to washing with PBS. Cells were permeabilized using 0.5 % Triton-X 100 in PBS for 15 minutes at room temperature and washed again with PBS. The copper-catalyzed click reaction occurred in 100 mM HEPES pH=8, 150 mM NaCl, 5 mM sodium ascorbate, 100 µM CuSO_4_, 500 µM THPTA and 20 µM sulfo-Cy5-azide for 30 minutes at room temperature. This and all following steps occurred under protection from light. The reaction solution was discarded and slides washed once with TBST (50 mM Tris-Cl,150 mM NaCl, 0.1 % Tween-20). HOECHST33342 was applied at a concentration of 10 µM in TBSP for 10 minutes, slides again washed twice with TBST and mounted using ProlongGold antifade mountant.

Imaging was performed on a Leica SP5 (Leica) using a 63x oil emersion objective. Detection of sulfo-Cy5-EU occurred using the default Leica Cy5 filter settings with excitation at 633 nm and detection at 650-750 nm wavelength.

### Immunoflourescence Staining and Confocal Microscopy

For the visualization of EXOSC2, MCF7 cells were grown on glass cover slips for 3 days. Sodium arsenite was applied at 400 µM for 0, 5 or 30 minutes, the media was discarded and cells washed once with PBS. Fixation occurred with 3 % paraformaldehyde in PBS at room temperature for 10 minutes subsequent to washing with PBS. Cells were permeabilized using 0.5 % Triton-X 100 in PBS for 15 minutes at room temperature and washed again with PBS. The antibody was diluted 1:100 in TBST (50 mM Tris-Cl, 150 mM NaCl, 0.1 % Tween-20) and binding allowed to occur over night at 4° C. Slides were washed twice with TBST before the secondary antibody (goat anti-mouse Cy5) was applied at a 1:500 dilution in TBST for 2 hours. Slides were stained with HOECHST33342 and imaged as described above.

### XRNAX-Assisted Crosslinking and Immunoprecipitation Followed by Sequencing (XRNAX-CLIP-seq)

For CLIP-seq from XRNAX extracts we first validated antibodies for normal IP by MS using the IP buffer 1 x (tris-Cl 50 mM pH=7.5, 0.5 % NP40, 150 mM LiCl, 0.1 % LiDS) later used for the CLIP-seq experiment.

For each sample XRNAX extracts were prepared as described above. For RNA fragmentation 10 mM tris-Cl and 5 mM EDTA were added to 100 µg XRNAX extract, which were sonicated in microTUBEs with AFA fiber (Covaris, 520045) using a S220 focused-ultrasonicator (Covaris) with the settings: 900 seconds, peak power 175, duty factor 50, cycles/burst 200 and average power 87.5.

For the size-matched input control (SMI-control) 5 µg of the sonicated XRNAX extract (approx. 2 µl) were mixed with 33 µl MilliQ water, 5 µl FastAP buffer 10 x and 10 µl FastAP. Dephosphorylation occurred for 15 minutes at 37 °C, then FastAP was inactivated for 5 minutes at 80 °C and the sample transfered to ice. 5 µl PNK buffer 10 x, 10 µl ATP 10 mM, 25 µl MilliQ and 10 PNK were added and incubated another 15 minutes at 37 °C. 15 µl of the SMI-control (approx. 200 ng RNA) was combined with 5 µl SDS-loading dye 5 x (tris-Cl pH=6.8 250 mM, SDS 10 %, 0.02 % bromphenol blue, glycerol 30 %) and 5 µl DTT 1 M. Samples were heated to 70 °C for 15 minutes before they were run on an SDS-PAGE along with the IP samples.

For the IP 100 µg sonicated XRNAX extract in approx. 125 µl was combined with 125 µl IP buffer 2 x (tris-Cl 100 mM pH=7.5, 1 % NP40, 300 mM LiCl, 0.2 % LiDS). 1 µg antibody was added for 4 hours at 4 °C on a rotating wheel before antibody capture with 100 µl protein G beads overnight.The beads were collected on a magnetic stand and the supernatant discarded. The beads were washed three times with 1 ml IP buffer, each time carefully turning the tube upside down until the beads were completely resuspended. Subsequently, beads were washed twice with 1 ml TBST while on the magnet. For end-repair the beads were resuspended in 100 µl dephosphorylation mix (80 µl MilliQ, 10 µl FastAP buffer 10 x, 8 µl FastAP, 2 µl RNASin) and incubated for 15 minutes at 37 °C, 1000 rpm shaking. Beads were collected on a magnetic stand, the supernatant discarded and the beads washed twice with 1 ml TBST while on the magnet. Subsequently, the beads were resuspended in 100 µl PNK mix (70 µl MilliQ water, 10 µl ATP 10 mM, 10 µl PNK buffer A 10 x, 8 µl PNK, 2 µl RNASin) and incubated for another 15 minutes at 37 °C, 1000 rpm shaking. Beads were collected on a magnetic stand and the supernatant discarded. Protein-RNA complexes were eluted into 5 µl SDS loading dye 5 x, 5 µl DTT 1 M and 15 µl MilliQ for 15 minutes at 70 °C. Beads were collected on a magnetic stand and the IP sample transferred to a fresh tube.

IP and SMI control were run alongside on a 4-12 % SDS-PAGE (NuPAGE BisTris, MES buffer) and blotted onto nitrocellulose with 500 mA for one hour at 4 °C. The area corresponding to the molecular weight of the protein of interest plus 75 kDA were excised, cut into pieces and transferred to a fresh tube. RNA was released by digestion with 50 µl proteinase K in 200 µl proteinase K buffer (tris-Cl 50 mM, EDTA 10 mM, NaCl 150 mM, SDS 1 %) at 55 °C for 30 minutes. 250 µl PCI for RNA was added, the sample mixed by inversion, incubated 10 minutes on ice and spun down 10 minutes with 12000 g at 4 °C. 200 µl of the aqueous phase were transferred to a fresh tube, NaCl added to a final concentration of 300 mM, combined with 1 µl GlycoBlue and 200 µl isopropanol. Samples were mixed and precipitated for 2 hours at −20 °C before centrifugation with 18000 g at −10 °C for 1 hour. Pellets were washed with 80 % ethanol and resuspended in nuclease-free water.

RNA produced by this protocol was approx. 30-80 nt in size, carried a 5’ phosphate and a 3’ hydroxyl. For generation of sequencing libraries we used the NextFlex Small RNA 3.0 kit and gel-based size-selection of RNA fragments from 30-50 nt according to the manufacturer’s instructions. Samples were barcoded so that twelve samples could be run in one lane on a HiSeq2000 (Illumina).

For a detailed version of this protocol and technical advice visit www.xrnax.com/applications.

### Processing and Analysis of XRNAX-CLIP-seq Data

Unique molecular identifiers (UMIs) were extracted using Je. Exact PCR duplicates were removed using BBMap. Adapters were trimmed using cutadapt and processed reads sequentially aligned to the 45S pre-ribosomal RNA (NR_046235.3), Repbase (Bao et al., 2015) and the hg38 genome with STAR, reusing reads which escaped previous alignment efforts. Non-exact PCR duplicates were removed with Je. The csaw library was used to calculate 20 nt coverage windows for the 45S rRNA for each individual time point. Subsequently, the DESeq2 library was used to calculate fold changes from this coverage between duplicates of the IP and duplicates of the SMI control. P-values were corrected for multiple testing with Benjamini-Hochberg.

### RNA Sequencing

For RNA sequencing, 10 µg RNA (as determined by NanoDrop UV-spectroscopy neglecting the protein content of samples) were digested for 30 minutes at 55 °C in proteinase K buffer (50 mM Tris-Cl, EDTA 5 mM, NaCl 150 mM, SDS 1%) using 10 µl proteinase K. Note that both TRIZOL and XRNAX extracted samples were treated identically. Subsequently, RNA was cleaned-up using the RNeasy mini kit (Qiagen) and was ready for RNA-Seq library preparation.

Specifically, for TRIZOL and XRNAX extracted RNA derived from MCF7 cells that were crosslinked at 254 nm wavelength (Figure 1C) RNA library preparation occurred with the TruSeq RNA Library Prep Kit v2 (Illumina, not stranded) after conditional depletion of ribosomal RNA (rRNA) using the Ribo-Zero rRNA Removal Kit (Illumina). Biological duplicates extracted with TRIZOL or XRNAX (4 samples in total) were barcoded to be sequenced in one lane on a HiSeq2000.

For TRIZOL and XRNAX-extracted RNA from 4SU-labeled MCF7 cells, library preparation occurred with the TruSeq Stranded Total RNA kit (Illumina, stranded) after depletion of rRNA using the Ribo-Zero rRNA Removal Kit. Biological duplicates extracted with TRIZOL or XRNAX (4 samples in total) were barcoded to be sequenced in one lane on a HiSeq2000.

### Processing and Analysis of RNA Sequencing Data

For the estimation of the rRNA content of libraries, which had not been RiboZero depleted, reads were aligned to a collection of human ribosomal sequences of the hg19 assembly retrieved using the UCSC table browser. The table used was ‘rmsk’ and filtering was applied so that ‘repClass does match rRNA’. All reads were aligned to those sequences using bowtie2 and reads that failed to align were written to a new file. Reads in this file were aligned to the complete hg19 assembly. Percentages of the rRNA content were estimated by comparing the number of reads aligning to hg19 rRNA sequences and residual reads aligning to the complete hg19 assembly.

For estimating the content of RNA biotypes in RiboZero depleted libraries, reads were aligned to the hg19 assembly using bowtie2. Subsequently, counting was performed with HTSeq-count using the geneset annotated by GENCODE 19 (release 12.2013) and using the GTF feature ‘gene’ for counting.

